# Constitutive Endolysosomal Perforation in Neurons allows Induction of α-Synuclein Aggregation by Internalized Pre-Formed Fibrils

**DOI:** 10.1101/2023.12.30.573738

**Authors:** Anwesha Sanyal, Gustavo Scanavachi, Elliott Somerville, Anand Saminathan, Athul Nair, Athanasios Oikonomou, Nikos S. Hatzakis, Tom Kirchhausen

**Affiliations:** Department of Cell Biology, Harvard Medical School, 200 Longwood Ave, Boston, MA 02115, USA; Program in Cellular and Molecular Medicine, Boston Children’s Hospital, 200 Longwood Ave, Boston, MA 02115, USA; Department of Chemistry University of Copenhagen, 2100 Copenhagen, Denmark; Department of Pediatrics, Harvard Medical School, 200 Longwood Ave, Boston, MA 02115, USA

**Author notes:** Corresponding author: Tom Kirchhausen. Ph.D. Harvard Medical School 200 Longwood Ave Boston, MA 02115 /.

**Keywords:** endolysosomal membrane traffic, live-cell imaging, PIKfyve kinase, Apilimod, Vacuolin-1, Parkinson’s disease, neurodegeneration

## Abstract

The endocytic pathway is both an essential route of molecular uptake in cells and a potential entry point for pathology-inducing cargo. The cell-to-cell spread of cytotoxic aggregates, such as those of α-synuclein (α-syn) in Parkinson’s Disease (PD), exemplifies this duality. Here we used a human iPSC-derived induced neuronal model (iNs) prone to death mediated by aggregation in late endosomes and lysosomes of endogenous α-syn, seeded by internalized pre-formed fibrils of α-syn (PFFs). This PFF-mediated death was not observed with parental iPSCs or other non-neuronal cells. Using live-cell optical microscopy to visualize the read out of biosensors reporting endo-lysosome wounding, we discovered that up to about 10% of late endosomes and lysosomes in iNs exhibited spontaneous constitutive perforations, regardless of the presence of internalized PFFs. This wounding, absent in parental iPSCs and non-neuronal cells, corresponded to partial damage by nanopores in the limiting membranes of a subset of endolysosomes directly observed by volumetric focused ion beam scanning electron microscopy (FIB-SEM) in iNs and in CA1 pyramidal neurons from mouse brain, and not found in iPSCs or in other non-neuronal cells in culture or in mouse liver and skin.

We suggest that the compromised limiting membranes in iNs and neurons in general are the primary conduit for cytosolic α-syn to access PFFs entrapped within endo-lysosomal lumens, initiating PFF-mediated α-syn aggregation. Significantly, eradicating the intrinsic endolysosomal perforations in iNs by inhibiting the endosomal Phosphatidylinositol-3-Phosphate/Phosphatidylinositol 5-Kinase (PIKfyve kinase) using Apilimod or Vacuolin-1 markedly reduced PFF-induced α-syn aggregation, despite PFFs continuing to enter the endolysosomal compartment. Crucially, this intervention also diminished iN death associated with PFF incubation.

Our results reveal the surprising presence of intrinsically perforated endo-lysosomes in neurons, underscoring their crucial early involvement in the genesis of toxic α-syn aggregates induced by internalized PFFs. This discovery offers a basis for employing PIKfyve kinase inhibition as a potential therapeutic strategy to counteract synucleinopathies.

## INTRODUCTION

The endocytic pathway is both an essential route of molecular uptake in cells and a potential entry point for pathology-inducing cargo. The cell-to-cell spread of cytotoxic aggregates, such as those of α-synuclein (α-syn) in Parkinson’s Disease (PD), exemplifies this duality. A hallmark of PD, which affects approximately 10 million individuals worldwide (Lashuel et al., 2013), are Lewy bodies, intracellular inclusions composed largely of α-syn aggregates and fragmented membranes, but α-syn also has a crucial role in normal synaptic function, probably by influencing synaptic vesicle trafficking (Iwai et al., 1995; Jakes et al., 1994; Courte et al., 2020).

α-syn, a 14 kDa neuronal protein, constitutes approximately 1% of cytosolic proteins in neurons (Stefanis, 2012). In its native state, α-syn is an unstructured monomer that can adopt an α-helical conformation upon interacting with phospholipids (Bartels et al., 2011; Wang et al., 2011; Burré et al., 2013; Breydo et al., 2012; Theillet et al., 2016; Rovere et al., 2018; Galvagnion et al., 2015; Weinreb et al., 1996; Eliezer et al., 2001; Kramer and Schulz-Schaeffer, 2007). Under pathological conditions, however, α-syn undergoes a conformational shift, forming cross β-sheet-rich amyloid fibrils that contribute to neurotoxicity (Conway et al., 2001). The transition of α-syn from a soluble, unfolded polypeptide to insoluble fibrillar aggregates is a critical aspect of PD pathogenesis. This process, known as α-syn aggregation, follows a nucleation-polymerization mechanism, in which pre-formed oligomers act as seeds for the assembly of fibrillar structures, resulting in the formation of Lewy bodies (Weinreb et al., 1996; Fauvet et al., 2012; Suzuki et al., 2018; Kramer and Schulz-Schaeffer, 2007; Desplats et al., 2009; Baba et al., 1998; Danzer et al., 2007).

*In vitro* studies using various central nervous system (CNS) derived models and *in vivo* experiments using brain tissues have contributed substantially to our understanding of α-syn pathology in PD (Luk et al., 2009; Volpicelli-Daley et al., 2011; Luna et al., 2018; Dryanovski et al., 2013; Masuda-Suzukake et al., 2013; Paumier et al., 2015; Luk et al., 2012; Desplats et al., 2009; Recasens et al., 2018; Nonaka et al., 2010; Redmann et al., 2017). These investigations have shown that α-syn aggregates disseminate in a ’prion-like‘ fashion, propagating from one cell to another. Thus, when neurons expressing wild-type (wt) α-syn are exposed to pre-formed fibrils (PFFs) of α-syn, these fibrils serve as a template or seed, catalyzing the intracellular formation of α-syn aggregates and damaging the affected neurons.

α-syn aggregates that form in cells upon incubation with PFFs localize predominantly within lysosomes (Bayati et al., 2022; Domert et al., 2016; Xie et al., 2022; Karpowicz et al., 2017; Konno et al., 2012). Freeman and colleagues (Freeman et al., 2013) proposed that this mechanism requires disruption of the lysosomal membrane, which they suggested would be caused by the internalized PFFs themselves. Others have proposed that in neuronal tissues, cell-to-cell propagation of α-syn aggregates might occur by transfer within lysosomes from donor cells to recipient cells through tunneling nanotubes (TNTs) (Senol et al., 2021; Abounit et al., 2016).

While PFF-mediated aggregation of α-syn is typically observed in neuronal cells, it can also occur in non-neuronal cells. For instance, human HEK293T cells ectopically expressing wild-type α-syn can form aggregates when PFFs are introduced by protein transfection, e.g. by Lipofectamine (Woerman et al., 2015). A genome-wide CRISPR interference screen in HEK293T cells showed that depletion of 1-phosphatidylinositol 3-phosphate 5-kinase (PIKfyve) reduces this α-syn aggregation (See et al., 2021). PIKfyve is a 240-kDa class III lipid kinase located on endosomal membranes, shown to have a critical function in endo-lysosomal trafficking and autophagy (Bissig et al., 2017; Rutherford et al., 2006; Lartigue et al., 2009; Sharma et al., 2019; Karabiyik et al., 2021; Kim et al., 2014). Inhibition of PIKfyve, either genetically or pharmacologically, depletes endo-lysosomal PI(5)P and PI(3,5)P2 phosphoinositides and leads to enlarged endo-lysosomes, disrupted fission, and impaired formation of autolysosomes (Karabiyik et al., 2021; Sharma et al., 2019; Choy et al., 2018; McCartney et al., 2014; Krishna et al., 2016; Leo et al., 2021). See and colleagues suggest that inhibiting transport of PFFs from endosomes to lysosomes by lowering PIKfyve activity would reduce lysosomal damage and hence reduce release of PFFs into the cytosol, thereby preventing seeding of cytosolic α-syn aggregates. Others have proposed that pharmacological inhibition of PIKfyve might instead reduce lysosomal PFF content by increased endo-lysosomal fusion with the plasma membrane and consequent exocytic release of PFFs into the medium (See et al., 2021).

In the work reported here, we re-examined these questions with an experimental approach that avoided non-physiological protein transfection. We used human induced pluripotent stem cell (iPSC)-derived neurons (iNs), in which α-syn aggregates originated from wild-type α-syn PFFs internalized from the medium. Formation of toxic α-syn aggregates was then restricted to naturally occurring lesions in late endosomes and lysosomes. We found that a small but significant proportion of late endosomes and lysosomes were inherently perforated (“leaky”) in iNs, even in the absence of exposure to PFFs; incubation with PFFs did not exacerbate this damage. We detected these intrinsically impaired late endosomes and lysosomes by live cell fluorescence microscopy, using biosensors designed to detect continuity with the cytosol. High-resolution, focused ion beam scanning electron microscopy (FIB-SEM) corroborated these observations, both in iNs and in CA1 pyramidal neurons from mouse brain. In contrast, endolysosomes in parental iPSCs and other non-neuronal cells, whether in tissue culture or in the liver, appeared structurally intact, in agreement with current understanding.

Acute pharmacological inhibition of PIKfyve activity in iNs with Apilimod or Vacuolin-1 alleviated the leaks we detected in late endosomes and lysosomes. This intervention not only prevented the PFF-mediated aggregation of cytosolic α-syn induced by PFF exposure but also greatly reduced the neurotoxicity and associated neuronal cell death induced by the resulting α-syn aggregates. Our findings--that a subset of endosomes and lysosomes are perforated in neuronal cells and that PIKfyve inhibitors reduce perforation and minimize PFF-induced cytotoxicity – suggest that targeting PIKfyve kinase activity might represent a viable therapeutic strategy for preventing and treating synucleinopathies such as PD. This approach would offer a way to mitigate the progression of these neurodegenerative disorders by addressing a fundamental aspect of their cellular pathology.

## RESULTS

### Incubation of iNs with PFFs causes formation of α-syn-YFP aggregates

We used a tissue-culture model resembling early stages in the synucleinopathy of Parkinson’s disease using human iPSC-derived neurons (iNs) in which we could induce formation of α-syn aggregates from ectopically expressed, cytosolic α-syn fused to YFP (a-syn-YFP) by incubating the cells for 3 days with PFFs (Gribaudo et al., 2019). We produced the PFFs *in vitro* from recombinant, wild type α-syn expressed in *E. Coli* (Volpicelli-Daley et al., 2014). The iNs were generated by differentiation from iPSCs in response to expression of the neuronal transcription factor NGN2 (Zhang et al., 2013; Lagomarsino et al., 2021). Inspection of the iNs 14-21 days after onset of differentiation showed appearance of neurites emanating from the soma and expression of the neuronal marker MAP2 (detected by immunofluorescence; Fig 1A); neurites and MAP2 were absent in the parental iPSCs (Fig. 1B).

**Figure 1.**
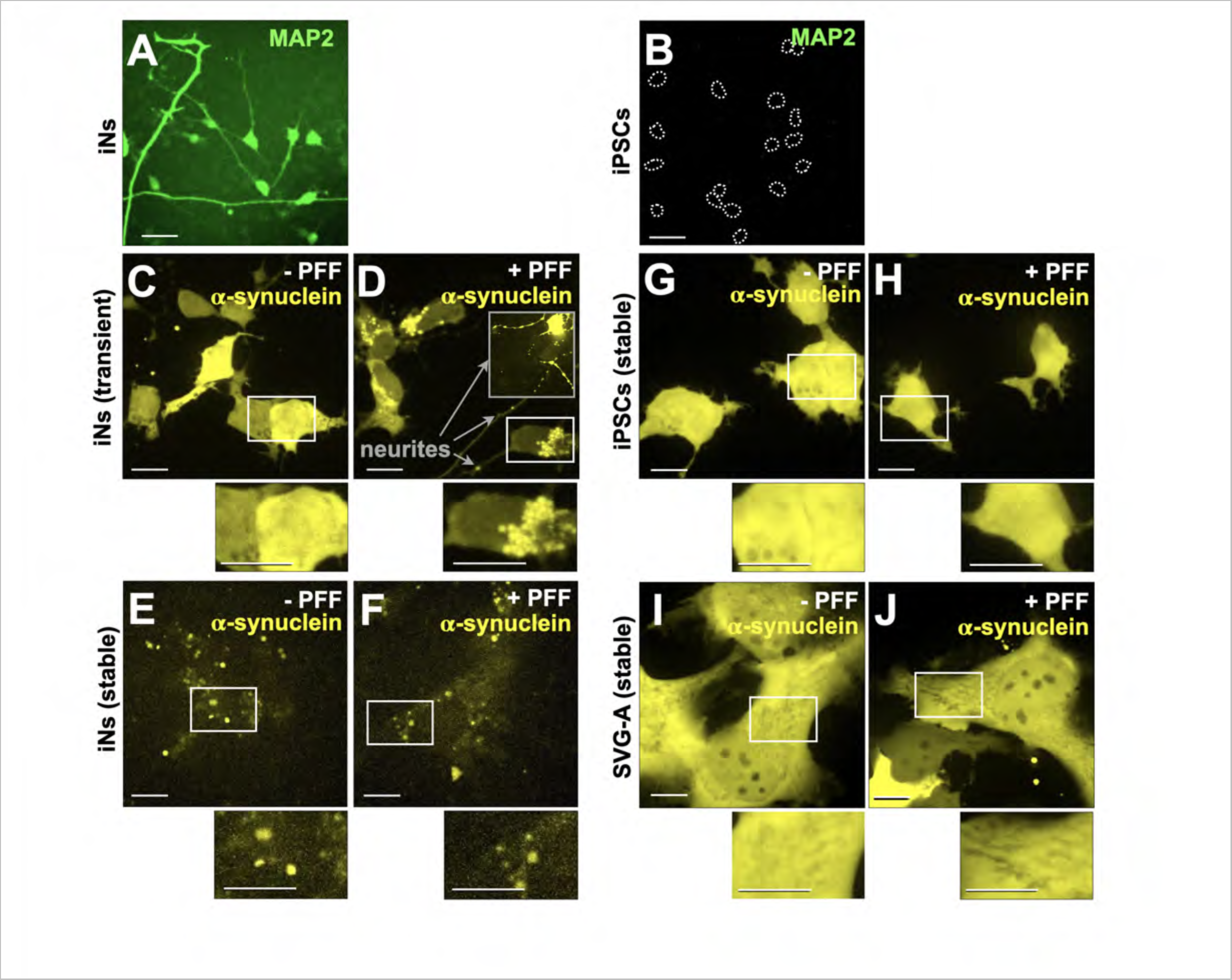
α-Synuclein-YFP Aggregation in iNs Exposed to Pre-formed Fibrils. **(A, B)** Immunofluorescence images depict **(A)** iPSC-derived neurons (iNs) and **(B)** iPSCs chemically fixed and treated with MAP2-specific antibody for neuronal identification. Cell perimeters in **(B)** are outlined with white dotted lines. Scale bar: 40 µm. **(C, D)** iNs with transient α-syn-YFP expression display **(C)** a diffuse cytoplasmic distribution without PFF treatment, compared to **(D)** distinct α-syn-YFP aggregates in soma and neurites after 3 days of incubation with 4 µg/ml PFFs. Scale bar: 10 µm. Insets provide 2x magnification, Scale bar: 5 µm. **(E, F)** iNs with stable α-syn-YFP expression, when incubated **(E)** without and **(F)** with 4 µg/ml PFFs for 3 days, show puncta consistent with α-syn-YFP aggregates. Scale bar: 10 µm. Insets offer 2x magnification, Scale bar: 5 µm. **(G, H)** iPSCs with stable α-syn-YFP expression exhibit a diffuse cytosolic pattern in the absence **(G)** or following a 3-day incubation with 4 µg/ml PFFs **(H)**. Scale bar: 10 µm. Insets show 2x magnification. Scale bar: 5 µm. **(I, J)** SVG-A cells stably expressing α-syn-YFP show a consistent diffuse cytoplasmic pattern both in the absence of **(I)** and after a 3-day incubation with 4 µg/ml PFFs **(J)**. Scale bar: 10 µm. Insets for 2x magnification, Scale bar: 5 µm.

Transient ectopic expression of α-syn-YFP in iNs yielded a diffuse, cytosolic fluorescent signal (Fig. 1C), a distribution expected for soluble cytosolic α-syn-YFP, also observed when stably expressed in immortalized astroglia-derived SVG-A cells (Fig. 1I), as in other non-neuronal cell types (Vasili et al., 2022; Furlong et al., 2000; Fortin et al., 2005; Imberdis et al., 2019). Continuous three-day incubation of iNs with PFFs added to the medium led to the appearance of α-syn-YFP fluorescent spots in both soma and neurites (Fig. 1D), consistent with the formation of α-syn-YFP aggregates, while only a cytosolic fluorescent α-syn-YFP signal was detected in similarly treated SVG-A cells (Figs. 1J). Equivalent experiments with undifferentiated parental iPSCs were not feasible, because iPSCs were not amenable to transient α-syn-YFP expression with our transfection protocol. Instead, using iPSCs harboring wild-type α-syn-YFP stably expressed by lentivirus transduction, we detected only diffuse cytosolic α-syn-YFP, regardless of whether or not they had been incubated for 3 days with PFFs (Fig. 1G, H). In contrast, iNs stably expressing α-syn-YFP imaged 5-14 days after the onset of differentiation displayed abundant intracellular α-syn-YFP spots and diffuse cytosolic α-syn-YFP, regardless of whether the iNs had been exposed to PFF for 3 days (Fig. 1E, F). We conclude that formation of α-syn-YFP aggregates induced by incubation with PFF is restricted to iNs and is not detected in cell types of non-neuronal origin. We note that transient or stable expression of ectopic α-syn-YFP in iPSCs or iNs was not toxic and did not induce cell death--an observation relevant for many of the following experiments.

### Incubation of iNs with PFFs mediates lysosomal aggregation of cytosolic α-syn-YFP

Published observations show that α-syn aggregates induced by internalized α-syn PFFs localize to LAMP1-containing lysosomes in human H4-neuroglioma derived cells (Jiang et al., 2017) mouse Cath.a-differentiated (CAD) cells (Senol et al., 2021) and mouse derived primary neurons (Volpicelli-Daley et al., 2011).

We used three different imaging fluorescence microscopy-based approaches to extend these observations and show that 3-day incubation of our iNs with PFFs also induced formation of α-syn-YFP aggregates localized to late endosomes and lysosomes. In the first approach (Fig. 2A-C, D-F), we used live cell spinning disc confocal fluorescence microscopy to visualize iNs transiently or stably expressing α-syn-YFP that had been incubated two hours before imaging with fluorescent Alexa Fluor 647 labeled Dextran (Dextran-AF647) an endocytic fluid phase marker known to accumulate in late endosomes and lysosomes and appear as intracellular fluorescent spots (Fig. 2A, B, D, E) (Ellinger et al., 1998). In the presence of PFF, these Dextran spots colocalized with most PFF-induced α-syn-YFP aggregates that formed in iNs transiently expressing α-syn-YFP (Fig. 2B, C). α-syn-YFP aggregates that spontaneously formed in iNs stably expressing α-syn-YFP and not exposed to PFFs failed to colocalize with internalized Dextran-AF647, suggesting in that case a cytosolic rather than endosomal location of the α-syn-YFP aggregates (Fig. 2D). In contrast, 3-day incubation with PFFs led to partial colocalization (Fig. 2E, F). These observations are consistent with endolysosomal localization of PFF-induced α-syn aggregates.

**Figure 2.**
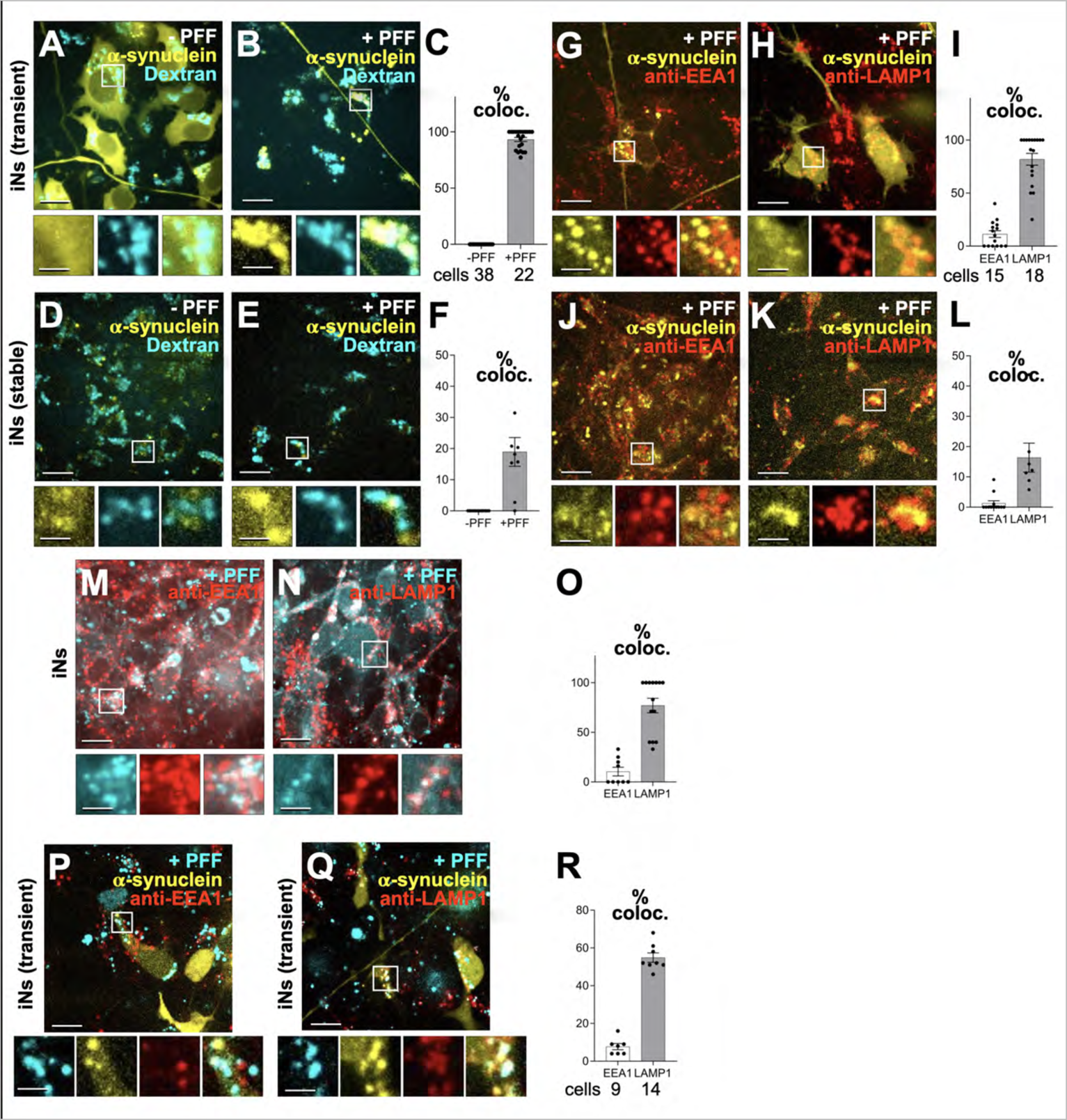
Localization of PFF-Induced α-Syn-YFP Aggregates in Endosomal and Lysosomal Compartments. **(A-C)** iNs transiently expressing α-syn-YFP incubated for 3 days with **(A)** 0 or **(B)** 4 µg/ml PFF, followed by a 2-hour incubation with 20 µg/ml fluorescent Dextran Alexa Fluor 647. Scale bar: 10 µm. Insets with 3x magnification, Scale bar: 3.3 µm. Histogram **(C)** quantifies the localization of α-syn-YFP and Dextran spots, indicating their predominant colocalization in late endosomes. Each dot represents data from all the spots within an individual cell; percentage of colocalization and cell count are noted. **(D-F)** iNs with stable α-syn-YFP expression incubated for 3 days with **(D)** 0 or **(E)** 4 µg/ml PFF, followed by 20 µg/ml Dextran Alexa Fluor 647 incubation. Scale bar: 10 µm. Insets with 3x magnification. Scale bar: 3.3 µm. Histogram **(F)** shows minimal colocalization between α-syn-YFP aggregates formed in the absence of PFF and Dextran, suggesting significantly lower endosomal location of the spontaneously formed α-syn-YFP aggregates. Each dot represents data for all the spots within a given field with an estimate of 5-9 cells; cells could not be defined because of weak cytosolic signal of α-syn-YFP. Colocalization percentage is specified. **(G-L)** iNs expressing α-syn-YFP transiently **(G, H)** or stably **(J, K)**, incubated with 4 µg/ml PFF for 3 days, were stained for EEA1 (early endosome marker, **G, J**) and LAMP1 (late endosome/lysosome marker, **H, K**). Scale bar: 10 µm. Enlargements with 3x magnification. Scale bar: 3.3 µm. Histograms **(I, L)** quantify colocalization of α-syn-YFP with EEA1 or LAMP1. Each dot in **(I)** represents integrated data from all the spots within a cell; colocalization percentages and cell counts are indicated. In **(L)**, each dot represents data for all the spots within a given field with an estimate of 5-9 cells; cells could not be defined because of weak cytosolic signal of α-syn-YFP. Colocalization percentage is specified. **(M-O)** iNs incubated with 4 µg/ml PFF-AF647 for 3 days and immuno stained for EEA1 **(M)** or LAMP1 **(N)**. Scale bar: 10 µm. Enlargements with 3x magnification, Scale bar: 3.3 µm. Histogram **(O)** quantifies colocalization between PFF-AF647 and these markers. Each dot represents data from all the spots within a field with 5-9 cells; colocalization percentages are provided. **(P-R)** iNs transiently expressing α-syn-YFP incubated with 4 µg/ml PFF-AF647 for 3 days, stained for EEA1 **(P)** or LAMP1 **(Q)**. Scale bar: 10 µm. Insets with 3x magnification. Scale bar: 3.3 µm. Histogram **(R)** quantifies colocalization between α-syn-YFP aggregates associated with internalized PFF-647, and these markers. Each dot represents a cell’s colocalization data from all spots; total cell count is noted.

In the second approach (Fig. G-I, J-L), we chemically fixed iNs after the 3-day PFF incubation and before imaging by spinning disc confocal fluorescence microscopy, to determine the extent of colocalization of PFF-induced α-syn-YFP aggregates with antibody markers specific for early endosomes (EEA1) and lysosomes (LAMP1) (Humphries et al., 2011). In agreement with recent observations (Senol et al., 2021; Bayati et al., 2022), we found minimal colocalization of PFF-induced α-syn-YFP aggregates with EEA1 (Fig 2G, I) and extensive colocalization with LAMP1 (Fig. 2H, I). Three-day incubation with PFF of iNs stably expressing α-syn-YFP failed to colocalize the induced α-syn-YFP aggregates with EEA1 (Fig. 2J, L) and led to the appearance of only a small fraction of α-syn-YFP aggregates colocalizing with LAMP1 (Figs. 2K, L).

In the third approach (Fig. 2M-R), we incubated iNs expressing endogenous α-syn alone (Fig. 2M-O) or together with transiently expressed α-syn-YFP (Fig. 2P-R), with PFF fluorescently tagged with Alexa Fluor 647 (PFF-AF647) for 3 days followed by chemical fixation. We then used spinning disc confocal microscopy to determine the extent of colocalization of the internalized PFFs with early or late endosomes and lysosomes identified with antibodies specific for EEA1 or LAMP1, respectively. We again found minimal colocalization of internalized PFFs with EEA1 (Figs. 2 M, O and P, R) and extensive colocalization with LAMP1 (Figs. 2N, O and Q, R), indicating that transient expression of α-syn-YFP had no detectable effect in the endo-lysosomal traffic of internalized PFFs.

These colocalization observations, which were consistent with published results obtained with neuronal cells (Jiang et al., 2017; Volpicelli-Daley et al., 2011; Senol et al., 2021; Karpowicz et al., 2017; Konno et al., 2012), validate our use of iNs as a convenient model system to study the early formation of α-syn aggregates in endo-lysosomes following PFF internalization.

### Constitutive perforation of limiting membranes in late endosomes and lysosomes of iNs

Aggregation of α-syn within endo-lysosomes might be due to a breach in the integrity of the limiting membrane surrounding them (Senol et al., 2021). According to this model, cytosolic α-syn could reach internalized PFFs retained in the lumen of damaged endo-lysosomes, thus becoming seeds for the generation of PFF-mediated α-syn endo-lysosomal aggregates.

The limiting membrane of endo-lysosomes is a natural barrier that would ordinarily prevent exchange of macromolecules between lumen and cytosol. Endolysomal limiting membrNE damage exposes luminal oligosaccharides that recruit galectin-3 and-8, which in turn initiate repair mechanisms (Radulovic et al., 2018; Jia et al., 2020, 2018). Chimeras of a fluorescent protein with these galectins are used as biosensors to detect perforated or leaky endolysosomes; the endolysosomal membrane perforation causes their fluorescence to redistribute from a diffuse cytosolic signal and to appear as distinct fluorescent spots associated with the damaged organelle (Jia et al., 2020; Thurston et al., 2012) We tested the iNs generated as described above and made the unexpected observation that the iNs had numerous intracellular fluorescent spots of mCherry-galectin-3 (Fig. 3B) and mCherry-galectin-8 (Fig. 3D), consistent with loss of endo-lysosomal integrity, even though these cells had not been exposed to PFFs. We likewise observed distinct spots of mCherry-galectin-3 or mCherry-galectin-8 in iNs transiently co-expressing α-syn-YFP but not exposed to PFF (Fig. 3E, H). We ruled out the possibility that these galectin spots might have represented mCherry-galectin-3 or mCherry-galectin-8 internalized from the medium due to secretion or release from iNs expressing the chimeras, since the spots were only detected in the galectin-3 or galectin-8 expressor cells, but never in adjacent non-expressor cells. The parental iPSCs showed diffusely distributed cytosolic mCherry-galectin-3 (Fig 3A) and few mCherry-galectin-8 spots (Fig 3C), as generally observed in non-neuronal cells with intact endosomes and lysosomes (Freeman et al., 2013; Paz et al., 2010; Aits et al., 2015).

**Figure 3.**
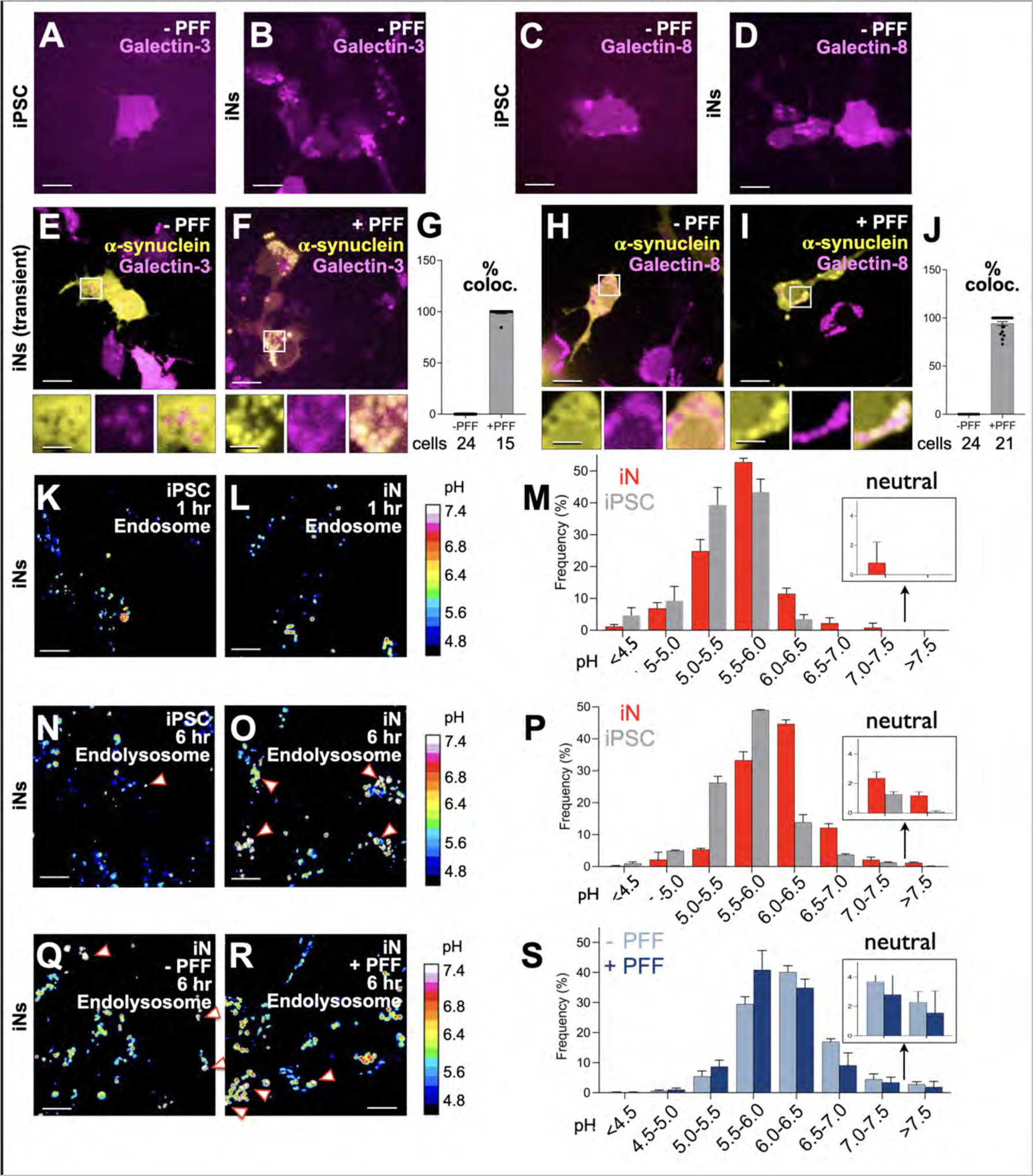
PFF-induced α-Syn-YFP Aggregates in perforated Endosomes and lysosomes. **(A, B)** iPSCs and iNs stably expressing mCherry-galectin-3, not exposed to PFFs, show diffuse cytosolic and punctate galectin signals, respectively. Scale bar: 10 µm. **(C, D)** iPSCs and iNs with stable mCherry-galectin-8 expression, unexposed to PFFs, display similar diffuse and punctate galectin patterns. Scale bar: 10 µm. **(E-J)** iNs transiently expressing α-syn-YFP with stable mCherry-galectin-3 **(E, F)** or mCherry-galectin-8 **(H, I)** expression, incubated for 3 days with **(E, H)** 0 or **(F, I)** 4 µg/ml PFF. α-Syn aggregates formed in PFF-treated cells colocalize with galectin-3 or galectin-8 puncta. Scale bar: 10 µm. Insets with 3x magnification. Scale bar: 3.3 µm. Histograms **(G, J)** show extent of colocalization between α-syn-YFP and mCherry-galectin-3 or α-syn-YFP and galectin-8 in iNs. Each dot represents data from all spots within a cell; colocalization percentages and cell counts are provided. **(K-P)** Live imaging of iPSCs and iNs incubated with 20 µg/ml each of pH-sensitive pHrodo Dextran 488 and pH-insensitive Dextran Alexa Fluor 560 for 1 hour **(K, L)** or 2 hours followed by a 4-hour chase **(N, O)**, labeling early/late endosomes and lysosomes, respectively. Scale bar: 10 µm. Heat map bars show pH values derived from fluorescence ratios in permeabilized samples incubated with solutions of different pH. Histograms **(M, P)** present average pH values ± standard deviation from dextran-containing spots, based on data from the 5366 and 10922 spots across 20 fields from iPSCs and 20 fields from three independently differentiated iN samples, respectively. **(Q-S)** iNs incubated with or without 4 µg/ml PFF for 3 days, then treated with the dextran mixture for 2 hours and a 4-hour chase. Heat map and histogram **(S)** as in **(K-P)**, with data from 3497 and 2239 spots across 10 fields from three independently differentiated iN samples.

While these observations were consistent with the existence of constitutively perforated, leaky or damaged endo-lysosomes, we also considered the possibility that repair of a transient opening had entrapped cytosolic galectins within the lumen of the now fully sealed compartment. To rule out this repair possibility, we monitored *in vivo* the pH of endosomes and lysosomes. We expected that only during opening, would the membrane rupture allow for the ionic equilibration with the cytosol required to neutralize the ordinarily acidic pH of the endo-lysosomal lumen (Maxson and Grinstein, 2014). To follow the pH, we used live cell, spinning disc, ratiometric fluorescence microscopy, implemented for *in vivo* pH imaging by incubating the iNs with a mixture of dextran tagged with pHrodo^TM^ Green (a pH sensitive fluorophore whose pKa ∼7.3 is best suited to quantify in the neutral pH range) and dextran tagged with pH-insensitive Alexa Fluor 560 (for content normalization). Fluorescence from the internalized mixture was monitored ratiometrically after one hr. of uptake or 4 hrs. after a 2-hr. period of dextran internalization, to deliver the label primarily into early-to-late endosomes (1 hr. time point) or into late endosomes-to-lysosomes (6 hr. time point), respectively (Fig. 3K-S). We found in iNs that neutral endosomes accounted for only ∼0.8 % of all labeled compartments at the early time point (Fig. 3 L, M) but for nearly 3.6 % at the later time point (Fig. 3 O, P). In contrast, the lumen of essentially all early/late endosomes and lysosomes in parental iPSCs remained acidic (Fig. 3K, M, N and P), as similarly observed with non-neuronal SVG-A cells. Our results also showed that late endosomes and lysosomes in iNs tended to be less acidic than their counterparts in iPSCs (Figs. 3M, P).

### Electron microscopic visualization of perforated limiting membranes in endosomes and lysosomes of iNs and neurons

What sort of endosomal or lysosomal opening accounts for loss of the usual pH gradient and for the local clustering of galectin-3 or-8. We used volumetric focused ion beam scanning electron microscopy (FIB-SEM) at approximately 5 nm isotropic resolution to examine iNs prepared by high pressure freezing and freeze substitution (HPFS), an approach that minimizes membrane perturbations (Studer et al., 2008). We could identify endosomes and lysosomes by their distinctive shape, size, and appearance. Unlike traditional single-plane imaging, our volumetric data allowed us to detect even small orifices at nearly any orientation with respect to the beam direction. We could indeed find many “open” endolysosomes (e.g., organelles containing a mixture of intraluminal vesicles and membrane fragments), usually with just a single nanoscale rupture of variable shape and typically between 100 and 300 nm in diameter, as depicted in the representative examples in Figs. 4 A, B. The staining pattern at the damage sites suggested minor leakage of luminal content into the cytosol.

**Figure 4.**
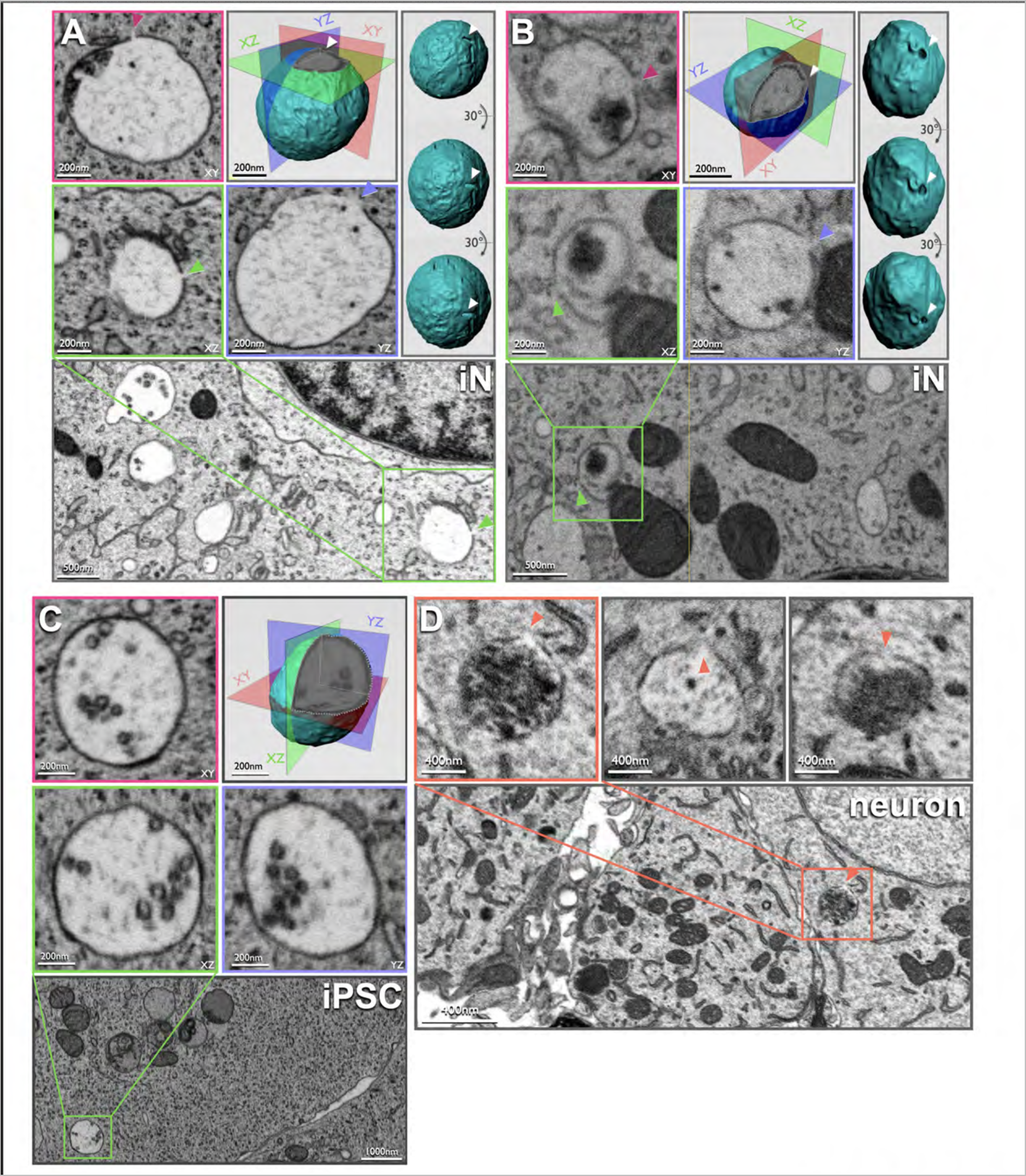
Visualization of perforated Endolysosomes in iNs and Rat Brain Neurons via FIB-SEM. **(A-C)** Orthogonal views, surface rendition and single plane view of FIB-SEM images acquired at 5 x 5 x 5 nm resolution of cells prepared by high pressure freezing and freeze substitution. Scale bars, 200 nm, 500 nm, 1000 nm. **(A, B**) highlight representative examples from two different iNs showing damaged endo-lysosomes with a small nanopore in their limiting membranes. **(C)** shows a representative example from a parental iPSC showing an intact endo-lysosome and lacking nanopores. **(D)** Top three panels show single plane views from three representative perforated endo-lysosomes imaged by volumetric FIB-SEM at 8 x 8 x 20 nm resolution from three different mouse brain CA1 pyramidal neurons prepared by chemical fixation (Sheu et al., 2022). Bottom panel shows an single plane lower scale view that includes the upper left panel. Scale bars, 400 nm, 4000 nm.

In contrast, endosomes, and lysosomes in parental iPSCs, as shown in Figure 4 C, and in non-neuronal cell lines such as SVG-A human fetal glial derived cells, HEK293A human epithelial derived cells, SUM159 human breast carcinoma derived cells, BSC-1 African green monkey kidney epithelial derived cells, U2OS human sarcoma derived cells (Gallusser et al., 2022) and HeLa human derived cells (Heinrich et al., 2021) showed no such openings, in agreement with the common view in the existing literature. Consequently, we propose that the membrane breaches in iNs directly observed using FIB-SEM represent the endo-lysosomal damage and the inferred interaction between luminal contents and adjacent cytosol we postulated from optical microscopy.

We further expanded our 3D visualization to neurons within an adult mouse brain, focusing on their endo-lysosomal integrity. Specifically, hippocampal CA1 pyramidal neurons in mature mouse brains were chemically fixed and visualized in our laboratory with volumetric FIB-SEM imaging at a resolution of 8 x 8 x 20 nm in our laboratory (Figure 4 D) (Sheu et al., 2022); we also analyzed similar images collected with 5.5 x 5.5 x 15 nm resolution at Janelia Research Campus (Sheu et al., 2022). The resulting data showed that these authentic, brain neurons contained a subset of endosomes and lysosomes with compromised limiting membranes, just as in our cell-culture model. By contrast, data from mouse liver (Parlakgül et al., 2022) and P7 mouse skin cells (OpenOrganelle, HMMI), for which imaging was conducted at 8 x 8 x 8 nm resolution, failed to display damaged endosomes, endo-lysosomes or lysosomes. Thus, there appears to be a neuron-specific propensity for a perforation to develop in the endo-lysosomal compartment.

### Pharmacological inhibition of endosomal PIKfyve kinase activity in iNs prevents PFF-induced α-syn-YFP aggregation

The phosphoinositides PI(5)P and PI(3,5)P2, generated by the endo-lysosomal phosphatidylinositol-3-phosphate/phosphatidylinositol 5-kinase (PIKfyve kinase), are essential for vesicular cargo traffic from late endosomes to lysosomes and for autophagosome maturation (Kim et al., 2014; Bissig et al., 2017; Rutherford et al., 2006; Choy et al., 2018; Sharma et al., 2019). Without these lipids, endosomes enlarge into distended, vacuole-like structures (Krishna et al., 2016; Bissig et al., 2017; Choy et al., 2018; Kang et al., 2020).

Acute PIKfyve kinase inhibition with Apilimod or Vacuolin-1 led to the expected enlarged endosomes in iNs (Figs 5-7) just as seen in non-neuronal cells (Kang et al., 2020; Cerny et al., 2004). This treatment did not hinder receptor-mediated endocytosis of transferrin-646 (Fig 5A, B, G; 5’ Tf pulse) or fluid phase uptake of Dextran-AF647 (Fig 5C, D, H; 3 hr. uptake) or PFF-AF647 (Fig. 5E, F, I; 3-day uptake). While iNs transiently expressing α-syn-YFP incubated for 3 days with PFFs generated PFF-mediated α-syn-YFP aggregates in endosomes and lysosomes (Figs. 6B, E, G, 7B), the aggregates failed to form in the absence of PFFs (Fig. 6C) or if Vacuolin-1 (Fig. 6F, H) or Apilimod (Figs. 6I-P, 7C) were present during PFF incubation. Treatment of iNs with Vacuolin-1 or Apilimod also significantly reduced the number of mCherry-galectin-3 (Fig. 6F, I-L) or mCherry-galectin-8 (Fig 6D, H, M-P) fluorescent spots, consistent with a direct correlation between inhibition of PIKfyve activity and loss of perforated endo-lysosomes. In contrast to iNs transiently expressing α-syn-YFP, iNs stably expressing α-syn-YFP and treated with Apilimod had α-syn-YFP aggregates that did not colocalize with internalized Dextran (Fig. 6S). Thus, Apilimod has no effect in these cells on aggregate formation in the cytosol, which is independent of PFF incubation (Fig. 6Q-S).

**Figure 5.**
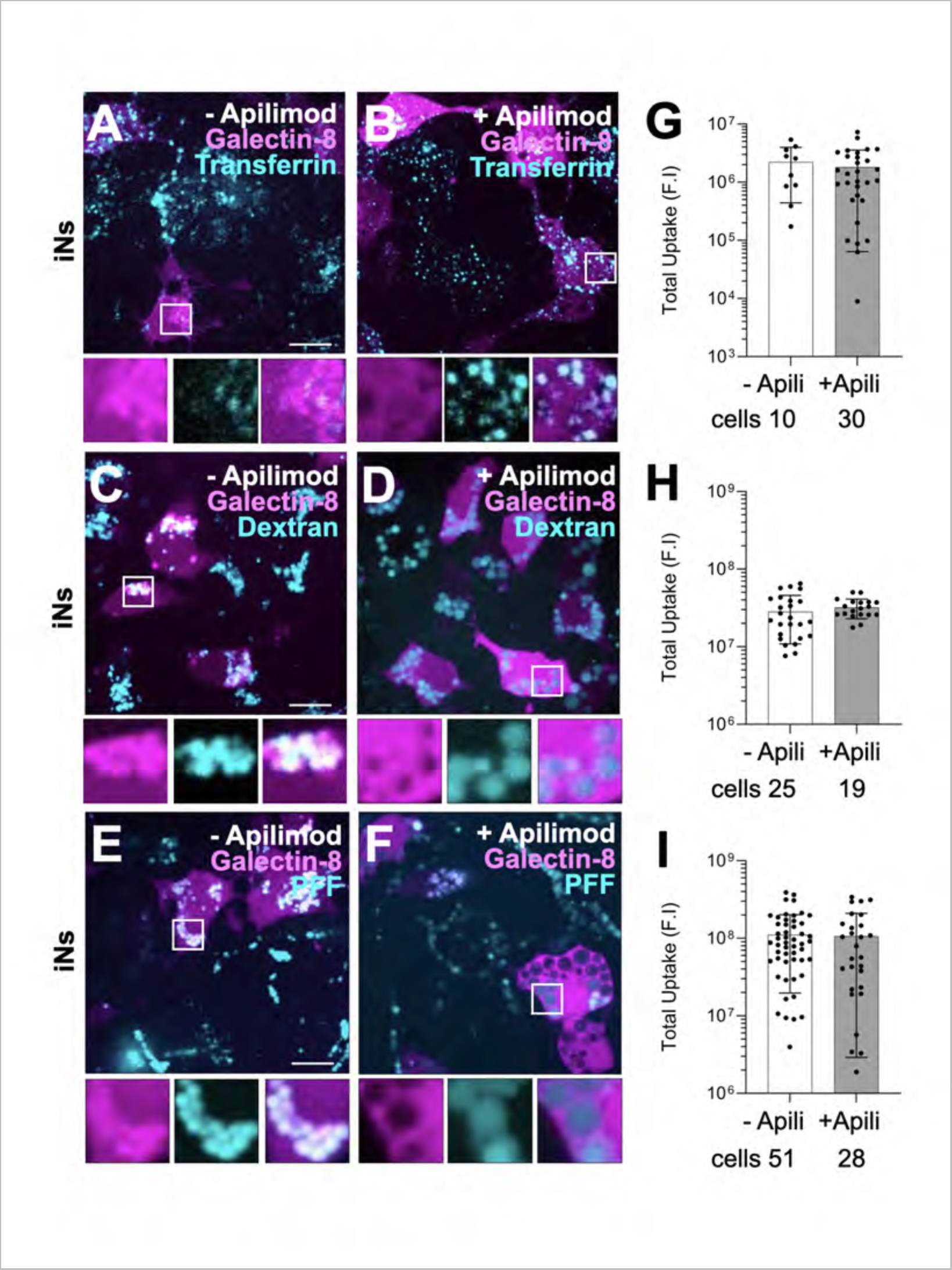
PIKfyve Kinase Inhibition reduces Endolysosomal Damage without Affecting Endocytosis. **(A-F)** iNs stably expressing mCherry-galectin-8 imaged using live 3D spinning disc confocal microscopy. Cells were incubated for 5 min. with 20 µg/ml Transferrin AF647 **(A, B)**, for 3 hrs. with 20 µg/ml Dextran AF647 **(C, D)**, or for 3 days with 4 µg/ml PFF-AF647 **(E, F)** in the absence **(A, C, E)** or presence **(B, D, F)** of 100 nM Apilimod. Notably, cells treated with Apilimod showed enlarged endosomes and lysosomes and did not exhibit galectin spots. The representative images are Z-max projections. Scale bar: 10 µm. Insets show 3x magnification, Scale bar: 3.3 µm. **(G-I)** Quantitative analysis comparing the lack of effect of 100 nM Apilimod treatment on the endocytic uptakes of Transferrin, Dextran AF647, and PFF-647 in the experiments highlighted by the examples depicted in **(A-F**). This quantification was determined by measuring the total fluorescence intensity of each endocytic marker per cell, utilizing live cell 3D-spinning disc confocal microscopy. Each dot represents the cumulative fluorescence intensity per cell; Cells, number of cells analyzed.

**Figure 6.**
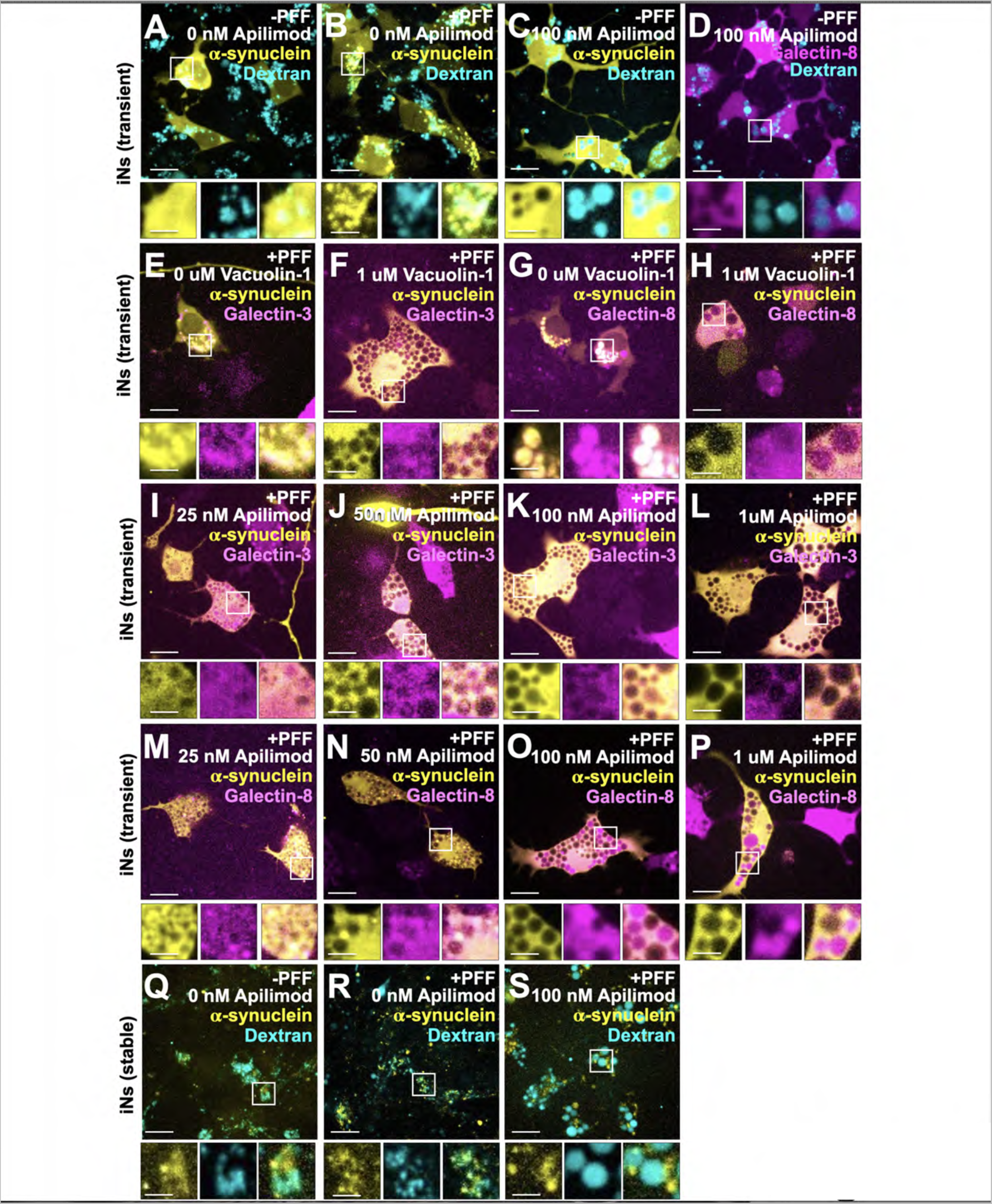
PIKfyve Kinase Inhibition prevents Endolysosomal Damage and PFF-Induced α-Synuclein Aggregation. **(A-P)** iNs transiently expressing α-syn-YFP and stably expressing either mCherry-galectin-3 or mCherry-galectin-8 were incubated for 3 days without, with 4 µg/ml PFF, and combinations of 4 µg/ml PFF with Apilimod or Vacuolin-1 at specified concentrations. A subset of cells **(A-C)** underwent a 2-hour treatment with Dextran AF647 prior to imaging. Scale bar: 10 µm. Insets show 3x magnification, Scale bar: 3.3 µm. **(Q-S)** iNs stably expressing α-syn-YFP were subjected to similar 3-day incubation protocols, including no treatment, exposure to 4 µg/ml PFF, and PFF combined with 100 nM Apilimod and including a 2-hr. pre-imaging uptake of Dextran AF647. Scale bar: 10 µm. Insets provide 3x magnification. Scale bar: 3.3 µm.

**Figure 7.**
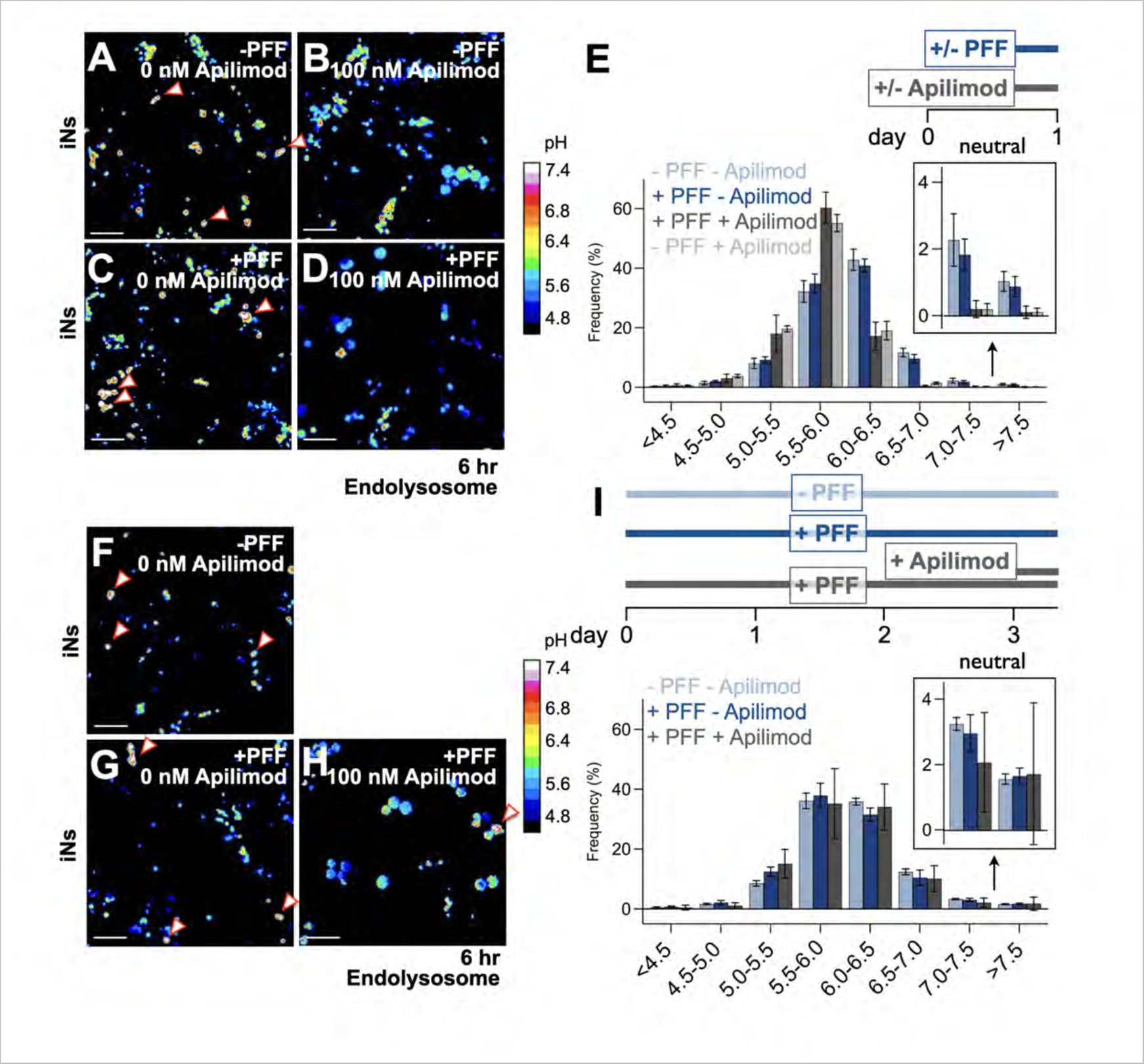
PIKfyve Kinase inhibition fails to rescue damaged Endolysosomal containing α-syn aggregates. **(A-D)** iNs without ectopic expression of α-syn-YFP were incubated for 8 hrs. with or without 4 µg/ml PFF, in the presence or absence of 100 nM Apilimod. Prior to *in vivo* pH imaging, the cells were incubated with a mixture of 20 µg/ml each of pH-sensitive pHrodo Dextran 488 and pH-insensitive Dextran Alexa Fluor 560; incubation was done for 2 hours followed by a 4-hour chase to label late endosomes, endolysosomes and lysosomes (*in vivo* pH imaging). Scale bar: 10 µm. Heat map bars show pH values derived from fluorescence ratios in permeabilized iNs incubated with solutions of different pH. **(D)** Histogram presents average pH values ± standard deviation from 8036, 5135, 2382 and 2575 dextran-containing spots across 6 fields from three independently differentiated iN samples treated with the indicated combinations of PFF and Apilimod. **(F-H)** Similar experimental design as in **(A-D)** except that the iNs were incubated with or without 4 µg/ml PFF for 3 days and 8 hrs., and the sample with PFF treated with 100nM Apilimod during the last 8 hours. These samples were then subjected to *in vivo* pH imaging as described for **(A-D)**. Scale bar: 10 µm. Heat map bars show pH values derived from fluorescence ratios in permeabilized iNs incubated with solutions of different pH. **(I)** Histogram presents average pH values ± standard deviation from 8896, 3838 and 271 dextran-containing spots across 6 fields from three independently differentiated iN samples treated with the indicated combinations of PFF and Apilimod.

In summary, acute pharmacological inhibition of PIKfyve kinase effectively mitigated constitutive endo-lysosomal damage in iNs and inhibited formation of PFF-mediated α-syn-YFP aggregates in endo-lysosomes, without impeding PFF uptake.

### Aggregation of cytosolic α-syn in iNs mediated by internalized PFFs occurs in constitutively perforated endolysosomes

We used the galectin localization assay described above to examine the correlation of endolysosomal perforation with α-syn-YFP aggregation. As expected, all the aggregates that appeared three days following 3-day exposure to PFFs co-localized with mCherry-galectin-3 (Fig. 3F, G) or mCherry-galectin-8 spots (Fig 3I, J). Despite PFF internalization, however, the overall number of galectin-positive sites appeared to be independent of whether PFFs were present in the medium or not. This result suggested that aggregate formation, which requires exposure of the internalized PFFs to cytosolic α-syn, might be occurring only in those endolysosomes with constitutive, PFF-independent perforation, rather than because of damage caused by the internalized PFFs, as previously proposed (See et al., 2021; Senol et al., 2021; Jiang et al., 2017; Freeman et al., 2013).

We therefore used the *in vivo* pH imaging to directly assess the association between endo-lysosomal perforation with internalized PFFs. We observed no alteration in the range or distribution of intraluminal pH values for late endosomes, endolysosomes and lysosomes in iNs after a 3-day PFF incubation (Fig. 3 Q-S). We conclude that in our cell model, internalized PFFs do not induce endo-lysosomal damage and that α-syn aggregate formation is a result of the normal presence of some leaky endolysosomes in the differentiated iNs.

We also used the *in vivo* pH imaging to compare the impact of PIKfyve kinase inhibition on the prevalence of endolysosomes with constitutive perforations. Our results revealed that an 8-hour incubation of iNs with PFFs in the presence of Apilimod abolished the subset of endolysosomes exhibiting a neutral luminal pH observed in iNs irrespective of PFF exposure (Fig. 7A-E). Conversely, pre-treating iNs with PFFs for 3 days, sufficient for the formation of α-syn aggregates, and subsequently co-incubating with PFF and Apilimod for 8 hours, did not alter the neutral endolysosome fraction, consistently accounting for approximately 5% of the total, regardless of PFF exposure (Fig. 7F-I).

We conclude that in our cell model, internalized PFFs do not induce endo-lysosomal damage and that α-syn aggregate formation is a result of the normal presence of some leaky endolysosomes in the differentiated iNs.

### Pharmacological inhibition of PIKfyve kinase activity protects neurons from death caused by PFF-induced α-syn endosomal aggregation

Prolonged exposure of primary neurons (Volpicelli-Daley et al., 2011; Redmann et al., 2017) or rodent brains (Desplats et al., 2009) to PFFs is toxic, ultimately leading to neuronal cell death. Using our iN-model system, we recapitulated similar toxicity in response to incubation with PFFs. Representative images of iNs incubated with cell impermeant BOBO-3 (to detect nucleic acids in dead cells) and cell permeant calcein AM (to detect viable cells) (Fig. 8) and the corresponding quantitative analysis (Fig. 9) showed a significant increase in the incidence of cell death when iNs were co-incubated with PFFs. Ten-day exposure to PFFs led to the accumulation of α-syn-YFP aggregates (Figs. 8A, B) and to the progressive death of up to ∼ 50% of the iNs, compared to only 10% of iNs not exposed to PFFs during the same period (Fig 9A-C, E). The extent of cell death was similar regardless of the absence (Fig. 9A) or presence of stably (Fig. 9B) or transiently (Fig. 9C, E) expressed α-syn-YFP.

**Figure 8.**
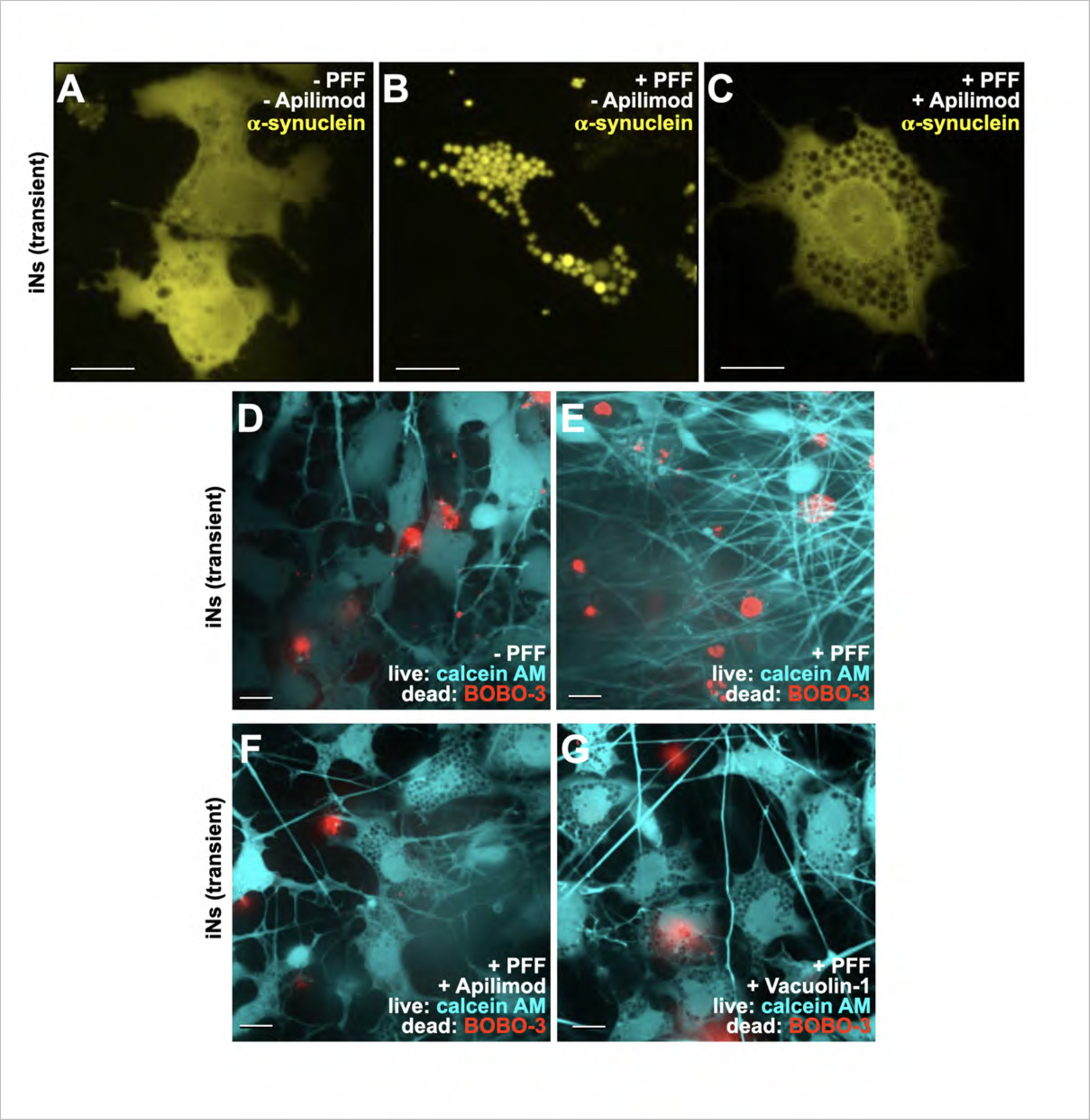
PIKfyve Kinase inhibition fails to rescue damaged Endolysosomal containing α-syn aggregates. **(A-C)** iNs transiently expressing α-syn-YFP were grown for 10 days in the absence or presence of 4 µg/ml PFF administration, with and without 100 nM Apilimod. Scale bar: 10 µm. **(D-G)** iNs with transient α-syn-YFP expression grown for 10-day with or without 4 µg/ml PFFs, alone and in conjunction with either 100 nM Apilimod or 1 µM Vacuolin. After incubation and prior to imaging, cells were incubated with calcein AM to identify live cells and BOBO-3-iodide for dead cells. Scale bar: 10 µm.

**Figure 9.**
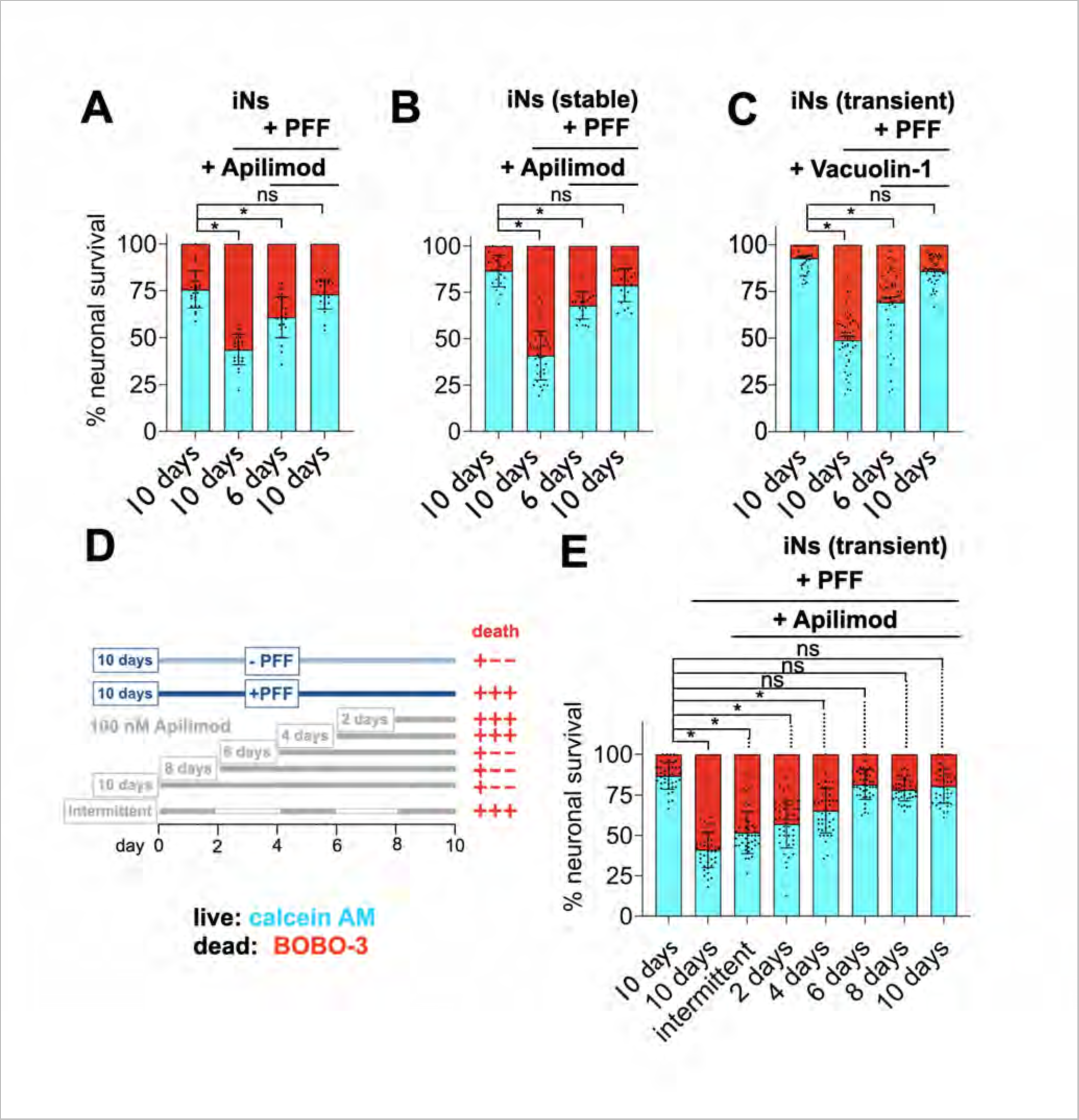
Quantitative Evaluation of Reducing PFF-Induced Neuronal Death by PIKfyve Kinase Inhibition. **(A)** The histograms summarize the quantification of live and dead cell in iNs under various experimental conditions, as detailed in Fig. 8. The data are presented for each imaging field, with values averaged and expressed as mean ± standard deviation. There were typically 45 fields, each with about 10-25 iNs. Statistical significance of the difference (p<0.0001) is indicated by a star symbol; ns means no statistical difference. **(B)** iNs not expressing α-syn-YFP were subjected to a 10-day incubation with or without 4 µg/ml PFF. Treatments with 100 nM Apilimod were either continuous throughout the 10 days or initiated during the final 6 days. Each point on the histograms represents the fraction of live or dead iNs cells within an imaging field. This condition mirrors **(A)**, but with iNs stably expressing α-syn-YFP. **(C)** Similar to **(A)** but conducted with iNs transiently expressing α-syn-YFP and treated with 1μM Vacuolin-1. **(D)** Schematic representation of the design and summary of results for the experiment depicted in **(E)**. **(E)** This setup is analogous to **(A)** and using iNs transiently expressing α-syn-YFP. In this case, 100 nM Apilimod was administered either throughout the entire 10-day period or restricted to the last 2, 4, 6, or 8 days. The intermittent schedule involved alternating 2-day incubation periods with Apilimod, starting from the first day of the 10-day duration.

Inhibition of PIKfyve kinase in iNs prevented PFF-mediated cell death. Representative images show absence of α-syn-YFP aggregates (Fig. 8C) and significantly reduced incidence of cell death when iNs were co-incubated for 10 days with PFF and Apilimod (Figs. 8D, E, F and quantification in Fig. 9A, B, D) or PFF and Vacuolin-1 (Fig. 8G and quantification in Fig. 9C). Comparable treatment during the last 6 days of PFF incubation afforded partial protection (Fig. 9A-C), but shorter Apilimod treatments in the last two or four days, or sequentially alternating two-day treatments on days 0, 4, and 8 did not (Fig. 9D). Off-target effects of Apilimod or Vacuolin-1 were unlikely, as Apilimod inhibited more strongly than Vacuolin-1, consistent with their relative inhibitory potencies for the enzymatic activity of PIKfyve kinase *in vitro* (Cai et al., 2014) and for the infectivity of PIKfyve-kinase dependent Zaire Ebola virus or SARS-CoV-2 infectivity *in vivo* (Kant et al., 2023; Kang et al., 2020). We conclude that pharmacological PIKfyve kinase inhibition by Apilimod or Vacuolin-1 treatment protects iNs from cell death by preventing endolysosomal damage rather than by directly reversing α-syn toxicity.

## DISCUSSION

The most striking finding from the results presented here is detection of a subset of constitutively leaky late endosomes and lysosomes in iNs and neurons in brain, due to nanoscale perforation of their limiting membranes. The membranes of endosomes and lysosomes in non-neuronal cells are all completely intact, as shown by many previous studies (Vest et al., 2022; Burbidge et al., 2022; Chou et al., 2023) as well as by experiments described here.

We have used several complementary approaches to gather evidence for constitutive endolysosomal damage in neurons, both in brain and iNs induced in the laboratory. We have visualized damaged structures directly by volumetric focused FIB-SEM and indirectly by fluorescence microscopy using probes for lysosomal pH (the lumen of endo-lysosomes will approach neutrality in the case of a perforated limiting membrane). We have also used recruitment of cytosolic, soluble mCherry-galectin-3 or mCherry-galectin-8 to detect leakage. We found galectin fluorescent spots in iNs, as others have done (Eapen et al., 2021), but very rarely in non-neuronal cells such as iPSCs and SVG-A cells, in accord with published reports on HEK292 cells (Chen et al., 2014) and fibroblasts (Vest et al., 2022). Moreover, work on human tNeurons transdifferentiated from primary fibroblasts and immuno-stained for endogenous Galectin-3 found galectin spots in the tNeurons but not in the parental fibroblasts (Chou et al., 2023). A similar pattern of galectin spots was seen in neurites from mouse hippocampal neurons transduced to express mCherry-Galectin-3 (Polanco et al., 2021), although not in analogous primary neurons also expressing mCherry-Galectin-3 (Calafate et al., 2016). Failure to find Galectin-3 spots in neuroblastoma-derived SH-SHY cells (Freeman et al., 2013; Burbidge et al., 2022; Flavin et al., 2017) could have been caused by incomplete differentiation at the time of imaging, possibly due to use of a suboptimal medium (Shipley et al., 2016). Although some of the galectin fluorescence we detected could have come from accumulation of re-internalized galectins secreted into the medium (Eapen et al., 2021), we believe that this potential additional source of fluorescent signal does not confound our conclusions, because they do not appear in non-expressor neighboring iNs.

The experiments here do not define the lifetime of the nanopores we detect, nor do they show whether only a specific endolysosomal subset can perforate or whether any late endosome or lysosome can acquire a potentially transient leak. Cells ordinarily avoid toxicity due to leakage from transient damage to the limiting membrane of compartments in the endolysosomal pathway by engaging the ESCRT-III mediated membrane repair mechanism (Skowyra et al., 2018; Jia et al., 2020; Radulovic et al., 2018). When extensive, the damage triggers sequestration of the affected organelles within autophagosomes, which are taken up and degraded by intact lysosomes, through a process termed “lysophagy” (Papadopoulos et al., 2017; Maejima et al., 2013). Sporadic instances of lysosomal damage occur in special circumstances, such as xenophagy--the selective degradation of damaged, bacteria-containing vacuoles or phagosomes (Mazin et al., 1987; Boyle and Randow, 2013). Similar processes have been detected in tissue samples from individuals with specific diseases--e.g., hyperuricemic nephropathy (Maejima et al., 2013; Emmerson et al., 1990) and inclusion body myopathy associated with frontotemporal dementia (Papadopoulos et al., 2017).

A second general finding from our work is that internalized PFFs remained in late endosomes and lysosomes and that the aggregation of host α-syn and the accompanying toxicity is likewise in close association with these subcellular compartments. The early induction of PFF-mediated aggregates at late endosomes and lysosomes appears to be exclusive to iNs, as we did not observe such aggregation in the parental iPSCs, nor did we find it in any other non-neuronal cells. Published accounts of PFF-induced α-syn aggregates within lysosomes include co-localization of ectopically expressed Galectin3-GFP in neuroblastoma-derived SH-SY5Y cells (Burbidge et al., 2022; Flavin et al., 2017; Freeman et al., 2013) and in murine catecholaminergic neuronal (CAD) cells from a transgenic mouse model (Senol et al., 2021). In the absence of PFFs in the medium, diffuse Galectin3 distribution in (presumably poorly differentiated--see above) neuroblastoma-derived SH-SHY cells (Burbidge et al., 2022; Flavin et al., 2017; Freeman et al., 2013) or in cells of essentially non-neuronal character like neuroglioma-derived H4/V1S-SV2 cells (Jiang et al., 2017) and CAD cells is consistent with the absence of leaky endolysosomes.

PFF-mediated α-syn aggregation throughout the cytosol occurs in cells, regardless of their origin, following introduction of PFFs (or other protein aggregates such as Tau fibrils) by transfection with lipofectamine (Trinkaus et al., 2021; Chen et al., 2019). This distinction between distributed aggregation induced by transfection and late endosomal or lysosomal aggregation induced by internalized PFFs introduced by incubation from the medium is important for understanding how such structures form and analyzing their consequences for cell physiology and cell viability. We have found spontaneous cytosolic α-syn aggregation in iNs stably expressing α-syn-YFP, even in the absence of PFFs in the medium, but not in the parental iPSCs or in neuroglioma-derived SVGA cells. These aggregates appear to be non-toxic, however. That is, PFF-induced aggregation within leaky late endosomes and lysosomes is toxic but spontaneous aggregation in the cytosol is not. We cannot at present provide an explanation for this distinction. Because cells (e.g., iNs, primary neurons) with constitutively leaky endosomes and lysosomes are healthy in the absence of endocytosed PFFs, any toxic stress signal from those endo-lysosomes must be specific for those that contain the α-syn aggregates. Although our data appear to rule out PFFs as initial agents of endolysosomal perforation, it remains possible that the larger aggregates they nucleate ultimately generate further damage.

A third important finding is that inhibition of PIKfyve kinase by Apilimod or Vacuolin-1 prevented PFF-induced endosomal and lysosomal α-syn aggregation, even though we showed that the PFFs had reached these subcellular compartments. These compounds also prevented PFF induced toxicity in iNs, although not if introduced after onset of aggregation. They cause endosomes and lysosomes to swell into spherical, vacuole-like, compartments and interfere with proper endolysosomal traffic. A proposed mechanism by which inhibition of PIKfyve kinase stimulates exocytosis and clearance into the medium of PFF-induced aggregates (Lee et al., 2013) appears to be inconsistent with our observation that incubating with Apilimod intermittently or during the last 2-4 days after initiation of the 10-day incubation period with PFFs failed to alleviate toxicity. We suggest instead that an important effect of inhibiting PIKfyve kinase is to prevent endolysosomal damage while allowing clearance of the organelles already damaged before adding the drug. This suggestion is consistent with the marked reduction in the number of constitutively leaky endo-lysosomes in iNs treated with Apilimod (or Vacuolin-1), even in the absence of PFF added to the medium.

Why do neurons have a subset of what appear to be constitutively leaky endo-lysosomes? One possibility is that their luminal content is viscous and while escaping to the cytosol it behaves as a local physical barrier to prevent the reseal of the damaged membrane. A second possibility is simply inefficiency in the membrane damage repair mechanism, which could also provide a functional signal to accelerate autophagic recycling in these cells, which cannot dilute toxic content by cell division. Although sorting out the complexities of this phenomenon are beyond the scope of the current study, we suggest that the constitutively perforated endolysosomal membranes detected in neurons could facilitate cytosolic access of endocytosed neurotoxic aggregates, including--in addition to α-syn--Huntingtin, Aβ, and tau. Substantiating this connection could advance understanding of the pathologies caused by these aggregates. Moreover, preventing lysosomal damage and hence toxicity by inhibiting PIKfyve kinase suggests a potential avenue for therapeutic intervention.

## MATERIALS AND METHODS

### Plasmids

We generated mCherry-Galectin3 and mCherry-Galectin8 constructs by subcloning from mCherry-Galectin8/pCDNA3.1 and mCherry-Galectin3/pCDNA3.1 into pENTR/TOPO vectors (Thermo Fisher K240020) through TOPO cloning. We then transferred these constructs into the pLX304 lentiviral vector via Gateway LR Clonase cloning according to the supplier’s guidelines (Thermo Fisher 11791100). Dr. Ulf Dettmer (Brigham and Women’s Hospital) provided the wild-type α-syn-eYFP/pCDNA3.1 construct, which we subcloned into pENTR/TOPO and subsequently into pLX301 using the same methods. We acquired wild-type α-syn/pET21a (Addgene 51486) courtesy of the Michael J. Fox Foundation.

### Inhibitors

Vacuolin-1 REF TK was custom synthesized; Apilimod (HY-14644) was purchased from MedChem Express.

### Cells

SVG-A cells were purchased from ATCC (CRL 8621) and cultured in MEM media (Corning 10-009-CV) supplemented with 10% FBS (Atlanta Biologicals S11150H). BR33 iPSCs, as described by (Paull et al., 2015; Lagomarsino et al., 2021), were kindly provided by Tracy Young Pearse of Brigham and Women’s Hospital. We cultured these cells in StemFlex media (Life Technologies A33493). For plating iPSCs and iNs, tissue culture plates were coated with Growth Factor Reduced Matrigel (Corning 354320) or Matrigel (Corning 354234), respectively, according to the following protocol: 0.5mg of Matrigel was resuspended in 5mL cold DMEM/F12 media and filtered through a 40μm cell strainer (Corning 352340). 6mL Matrigel was used to coat a 10cm tissue culture plate such that the final concentration of Matrigel is approximately 8.7μg/cm^2^.

For differentiation to induced neurons (iNs), we plated the iPSCs at a density of 100,000 cells/cm^3^and co-transduced them with lentiviruses carrying pTet-O-NGN2-puro and Fudelta GW-rtTA plasmids, following the methodology outlined by (Zhang et al., 2013). After two days of transduction, we continued to culture the cells for an additional seven days prior to cryopreserving them for future use.

We initiated differentiation by seeding thawed iPSCs on Matrigel coated plates at a density of 2 × 10^6 cells on 10 cm plates, using StemFlex media supplemented with 10 μM ROCK inhibitor (Stemcell Technologies 72304). We cultured the cells to 75% confluence. On day one, we replaced the medium with KnockOut media (Gibco 10829.018) enriched with KSR (KnockOut Serum Replacement, Invitrogen 10928-028), 1% MEM non-essential amino acids (Thermo Fisher 11140050), 1% GlutaMAX (Thermo Fisher 35050061), 0.1% BME (Invitrogen 21985-023), and 2 μg/mL doxycycline (Sigma D9891) to induce NGN2 expression. On day two, we changed the medium to a 1:1 mixture of KSR and N2B medium (DMEM F12, Thermo Fisher 11320033, with 1% GlutaMAX, 3% dextrose, N2-Supplement B, StemCell Technologies 07156, 5 μg/mL puromycin, Life Technologies A11138-03, and 2 μg/mL doxycycline). On day three, we replace the medium with N2B media supplemented with B27 (1:100; Life Technologies 17504-044), 5 μg/mL puromycin, and 2 μg/mL doxycycline. At day four, we froze the differentiated iNs in Neurobasal media (Gibco 21103-049) combined with B27 (1:50), 10 ng/mL BDNF (Peprotech, 450-02), 10 ng/mL CNTF (Peprotech, 450-13), 10 ng/mL GDNF (Peprotech, 450-10), 10 μM ROCK inhibitor, 5 μg/mL puromycin, and 2 μg/mL doxycycline, and 10% DMSO, termed NBM medium. We conducted experiments on iNs between days 11–21 post-thaw, cultivating them in NBM.

### Lentivirus preparation and transduction

We seeded HEK293T cells at 2.5 × 10^6 cells per 15 cm plate in DMEM (Thermo Fisher 11965118), enriched with 10% FBS (Atlanta Biologicals S11150H), 1X GlutaMAX (Thermo Fisher 35050061), 1X sodium pyruvate (Thermo Fisher 11360070), and 1X MEM non-essential amino acids (Thermo Fisher 11140050) to prepare HEKT media. On the next day, we transfected the cells with mCherry-Galectin 8/pLX304, mCherry-Galectin 3/pLX304, and wild-type α-syn/pLX301 plasmids using Lipofectamine 2000, following the manufacturer’s instructions (Thermo Fisher 12566014).

For the transfection, we mixed 24 μg of each lentiviral plasmid, 18 μg of psPAX2 gag/pol packaging plasmid (Addgene 12260), and 12 μg of pMD2.G VSV-G envelope plasmid (Addgene 12259) with 135 μL of Lipofectamine in 6.75 mL of OptiMEM media (Thermo Fisher 51985091). After a 20-minute room temperature incubation, we applied the transfection mixture dropwise to the HEK293T cells plated on 15 cm tissue culture dishes bathed with 14 mL of HEKT medium. Following a 6-hour incubation, we replenish with 14 mL of HEKT media and again after another 12 hours. We collected the virus-containing medium 8 hours later, centrifuged it at 500 x g for 5 minutes at 4°C to clear debris, and stored the supernatant in 1 mL aliquots at-80°C after flash-freezing in liquid nitrogen.

For transduction, we added 1 mL of the thawed virus-containing supernatant to iPSCs or SVGA cells (approximately 8 × 10^5), seeded 18 hours prior in a 6-well plate with 1 mL of mTESR1 (StemCell Technologies 85850) or MEM medium, respectively. We replaced the media with StemFlex media 12 hours post-transduction and added either 5 μg/mL Blasticidin or 5 μg/mL Puromycin, then cultured the cells for an additional 4 days. The surviving cells were grown in the same medium lacking antibiotics for 7 more days, after which the cells in the same medium supplemented with 10% DMSO were frozen overnight at-80°C using a Nalgene® Mr. Frosty freezing container and then kept in liquid nitrogen until future use.

### Transfection of iNs

The day following thawing, we plated 40,000 iNs into each well of an 8-chamber slide (Cellvis C8-1.5H-N). We then transfected them using Viafect (Promega E4981), adhering to the manufacturer’s instructions. For the transfection, we combined 0.2 μg of α-syn-eYFP/pCDNA3.1 plasmid with 0.6 μl of Viafect reagent and 100 μl of OptiMEM. This mixture was incubated at 37 °C for 20 minutes before being applied dropwise to the iNs bathed with 200 μl NBM media. After 24 hours, we replaced the cultures with 200 μl fresh NBM media. We conducted most experiments on these cells within 18-21-day post-differentiation; some experiments were started on day 11.

### Preparation of α-syn preformed fibrils (PFFs)

BL21(DE3) E. coli (New England Biolabs C2527H) underwent transformation with wild-type α-syn/pET21a. We selected single colonies to inoculate in Luria Broth (LB) containing 100 μg/mL ampicillin (P212121 GB-A-301-25). At an optical density (OD600) of ∼0.5, we induced cultures with 1 μM isopropyl 1-thio-β-d-galactopyranoside (IPTG, Sigma 6758) for 4 hours. Post-induction, we harvested the cells and resuspended the pellet in 5 mL of 20 mM Tris pH 8.0, 25 mM NaCl, followed by lysis via boiling for 15 min. We centrifuged the lysate at 20,000 x g for 20 minutes at 4 °C. We then applied the supernatant onto two tandem 5 mL HiTrap Q HP anion exchange columns (GE Healthcare) pre-equilibrated with 20 mM Tris pH 8.0, 25 mM NaCl. We eluted α-syn using a 5 mL linear gradient of 25–1000 mM NaCl in 20 mM Tris pH 8.0, 1 M NaCl. We pooled the α-syn-containing fractions and further purified the protein by size-exclusion chromatography on a HiPrep Sephacryl S-200 HR 16/60 gel filtration column (GE Healthcare) in 50 mM NH4Ac pH 7.4. We collected the peak fractions, aliquoted them into 20 μL volumes, lyophilized using a FreeZone-84°C lyophilizer (Labconco), and stored at −80 °C. Before use, a given aliquot was reconstituted by adding 5μl of 10 mM NH4Ac pH 7.4.

We followed this protocol to generate pre-formed fibrils (PFF): we used five reconstituted aliquots that were centrifuged at 21,130 × g for 20 minutes at 4°C. We kept the supernatant in a 1.5 mLmL Eppendorf tube and added 70-100 μL 1X PBS to achieve a 5 mg/mL protein concentration, confirmed using the BCA protein assay (Thermo Scientific A55864). After a brief vortex for 3 seconds at high speed, we placed the sample on an orbital shaker, set at 1,000 RPM and 37°C for 7 days, after which we placed the tube with the resultant PFFs in an ice bath and sonicated for 60 cycles (0.5 sec on, 0.5 sec off) at 10% amplitude using a QSonica XL-2000 sonicator with a 1/8” tip immersed in the solution. Subsequently, we aliquoted into 5 μL volumes, flash froze them in liquid nitrogen, and stored at-80°C until further use.

We prepared 4 mM stock solutions of Alexa 647 dye (Thermo Scientific A33084) by dissolving 1 mg in milliQ water, aliquoted in 10 μL volumes, dried using a speed vac (Savant), and stored at – 80°C. For fluorescently labeling PFFs, we thawed one aliquot, adjusted the protein concentration to 0.5 mg/mLmL by adding 85 mM NaH2CO3 (pH ∼8.3) and incubated in the dark at room temperature for 1 hour with Alexa647 at a final concentration of 0.05 mg/mL. To remove unbound dye after the labelling reaction, we utilized a Zeba 7K MWCO spin desalting column (Thermo Fisher 89882). The labeled protein was then aliquoted into 5 μL volumes, flash frozen in liquid nitrogen and stored at-80°C.

### Immunofluorescence

We began cell preparation by rinsing them with 1X PBS. We then fixed the cells with 4% (w/v) paraformaldehyde for 30 minutes. After discarding the fixative, we blocked the cells with 5% (v/v) BSA in 1X PBS for 30 minutes and permeabilized them with 0.1% (v/v) Triton X-100 in 1X PBS for 5 minutes at room temperature. Next, we incubated the cells with primary antibodies: MAP2 (1:1000; Abcam ab32454), LAMP1 (1:500; Abcam ab25630), or EEA1 (1:200; Santa Cruz sc-6415) diluted in 1X PBS for 1 hour at room temperature. We washed the cells thrice with 1X PBS for 10 minutes each. We then incubated the cells with Alexa Fluor-conjugated secondary antibodies for 1 hour at room temperature, followed by another three 10-minute washes with 1X PBS. Finally, we proceeded with immediate imaging in 1X PBS using spinning disk confocal microscopy or stored the cells at 4°C in 1X PBS for imaging on the following day.

### In vivo pH imaging

The pH of endosomes and lysosomes was estimated by incubating cells at 37°C for the indicated times with a mixture of 20μg/ml each of dextran tagged with pHrodo^TM^ Green (a pH sensitive fluorophore whose pKa ∼7.3 is best suited to quantify in the neutral pH range) and dextran tagged with pH-insensitive Alexa Fluor 560 (for content normalization). iNs or iPSCs were then washed thrice with 1X PBS, transferred to phenol-red free neurobasal medium supplemented with 1% B-27, 10 ng/mL BDNF, 10 ng/mL CNTF, and 10 ng/mL GDNF or Fluorobrite, respectively, and three biological replicates per experimental condition were ratiometrically imaged at 37°C and throughout the whole cell volume using spinning disc confocal microscopy.

The pH of the endosomes and lysosomes was estimated from pH calibration curves (Steinberg et al., 2010). Briefly, iNs or iPSCs cells were first incubated for 2 hrs. with 20μg/ml each of pHrodo Green-Dextran and Dextran-AF647, washed thrice in 1X PBS, followed by 15 min. incubation at room temperature with universal buffer (10 mM HEPES, 10 mM MES, 10 mM sodium acetate, 1 mM CaCl2, 140 mM KCl, 5 mM NaCl, and 1 mM MgCl2) and containing 1μM each of Nigericin (Cayman Chemicals 11437) and Monensin (Cayman Chemicals 16488). Samples were incubated with universal buffer titrated to different pH (4.4, 4.8, 5.2, 5.8, 6.0, 6.4, 6.8, 7.2, 7,4). Two biological replicate samples were volumetrically imaged for each pH using spinning disc confocal microscopy. Ratiometric quantification was done as described in Image Analysis. Calibration curves were generated using the sigmoidal dose-response function in GraphPad Prism 10.0.

### Cell survival assay

To determine the proportion of live and dead cells, we plated ∼ 40,000 iNs on Matrigel coated 8-chamber glass slide (Cellvis C8-1.5H-N) and cultured them for 11 days post-differentiation after which incubation with Apilimod and PFF were for 10 days as described elsewhere. Briefly, cells in 50 μL imaging media (phenol red-free Neurobasal medium supplemented with 1% B-27, 10 ng/mL BDNF, 10 ng/mL CNTF, and 10 ng/mL GDNF) were incubated for 15 minutes at room temperature with 50 μL 2x stock solution of LIVE/DEAD Cell Imaging Kit 488/570 (Thermo Scientific R37601) containing live cell Calcein AM and dead cell stain BOBO-3-Iodide, followed by spinning disc confocal imaging with 40x magnification and counting.

### Spinning Disk Confocal Imaging

We plated 40,000 cells per well on an 8-chamber slide (Cellvis C8-1.5H-N), pre-coated with Matrigel for iNs (grown up to 21 days before imaging) or GFR Matrigel for iPSCs (grown overnight before imaging). For iNs, we used phenol red-free Neurobasal medium (Thermo Fisher 12348017) supplemented with 1% B-27, 10 ng/mL BDNF, 10 ng/mL CNTF, and 10 ng/mL GDNF as the imaging medium. SVG-A cells were plated in chambers devoid of Matrigel. iPSCs and SVG-A cells were imaged in Fluorobrite medium, supplemented with 10% FBS and 25 mM HEPES pH 7.4. We mounted the slide chambers within a temperature-controlled, humidified chamber with 5% CO2 at 37°C for live cell imaging, while fixed samples were imaged at 25°C.

Images were acquired using spinning disk confocal microscopy controlled by Slidebook 6.4 software (3I) with the following configurations: a) A Marianas system (Intelligent Imaging Innovation) used for imaging of live or chemically fixed cells comprised a Zeiss Axio Observer Z1 microscope (Carl Zeiss) with a 20x (NA 0.5), 40x (NA 0.75) and 63× (NA 1.4) (Carl Zeiss) Apochromat objectives, a CSU-XI spinning disk unit (Yokogawa Electric Corporation), a heated stage (20/20 Technology), and a spherical aberration correction system (Infinity Photo-Optical). We utilized solid-state lasers operating at 405, 488, 561, and 640 nm (100 mW, Coherent Inc.) for excitation, modulated by an acoustic-optical tunable filter lasers linked to the spinning disc by a single mode fiber optic. Z-stacks were acquired at 270-nm z-intervals with 20-100 ms exposures using an air cooled QuantEM 512SC CCD camera (Photometrics). b) The same Marianas system with additional magnification of 1.2x, a heating stage (OKO Lab), a 3I spherical aberration correction system and a LaserStack^TM^ (3i) with diode lasers 405 (140 mW), 488 and 560 (150mW), 640 (100mW). Z-stacks were acquired at 270 nm intervals and 10 ms exposure using a sCMOS camera (Prim 95B, Teledyne Photometrics). c) a second Marianas system used to image chemically fixed cells was based on a Zeiss Axio Invert 200M microscope (Carl Zeiss) with a 63× Plan-Apochromat objective (NA 1.4, Carl Zeiss), a CSU-22 spinning disk (Yokogawa Electric Corporation), a heating stage (OKO Lab) and solid-state lasers operating at 491, 561, or 660 nm. Z-stacks were acquired at 270 nm intervals with exposure times between 50-100 ms using an air cooled QuantEM 512SC CCD camera (Photometric).

### High Pressure Freeze Substitution

We plated ∼200,000 cells on top of 6 x 0.1 mm sapphire disks (616-100; Technotrade International) coated with Matrigel for iNs and GFR Matrigel for iPSCs, themselves placed in 24 well dish containing 0.3 mL of appropriate medium, and cells allowed to grow 14 days for iNs and overnight for iPSCs at 37°C and 5% CO2. Cell viability (cell spread and shape) was verified by phase contrast using a tissue culture microscope. The sapphire disks, with cells adhered to one surface, were sandwiched between two aluminum platelets (Cavity 0.3 mm with one side ground 611; Technotrade International/Cavity 0.1/0.2 mm 610; Technotrade International). We subjected the cells to high-pressure freezing using a Wohlwend Compact 2 device (Technotrade, M. Wohlwend GmbH). The sapphire disks with adhered frozen cells were then immediately submerged in liquid nitrogen and placed into cryotubes on top of frozen freeze substitution solution (2% OsO4, 0.1% uranyl acetate, and 3% water in acetone).

Freeze substitution was carried out on an EM AFS1 automatic device (Leica Microsystems) according to the following schedule: a 2-hour hold at −140 to −90°C, a 24-hour hold at −90°C, a temperature ramp from −90 to 0°C during an elapse period of 12 hours, and finally, a 1-hour hold ramping from 0 to 22°C. We removed the cryotubes at room temperature and performed three sequential rinses with anhydrous acetone, propylene oxide (Electron Microscopy Sciences), and a 50% resin solution (24 g Embed 812, 9 g DDSA, 15 g NMA, 1.2 g BDMA; 14121; Electron Microscopy Sciences) in propylene oxide. We then transferred the samples to 100% resin placed within molds (EMS 70900). The resin-embedded samples were transferred to an oven at 65°C for 48 hours to polymerize the resin. To release the sapphire disks were released from the polymerized resin containing the cells using a thermal shock method, by immersing them first in liquid nitrogen and then in boiling water; the released resin block was then subjected to volumetric focused ion beam scanning electron microscopy (FIB-SEM).

### Focused Ion Beam Scanning Electron Microscopy

We prepared the resin blocks containing cells by sanding and then affixing them onto aluminum pin mount stubs (Ted Pella) with the cell-containing side exposed. We secured the blocks to the stubs using conductive silver epoxy adhesive (EPO-TEK H20S; Electron Microscopy Sciences), avoiding areas designated for imaging, and allowed the adhesive to cure for 24 hours at 65°C. Prior to FIB-SEM imaging, we coated the resin block surface with a 20 nm layer of carbon using a high-purity carbon cord source in a Quorum Q150R ES sputter coater (Quorum Technologies).

FIB-SEM imaging was done using a Zeiss crossbeam 540 microscope; we adjusted the stage for minimal eccentricity and tilt (54°), and working distance of 5 mm. Once a cell of interest was located by SEM, we further prepared the sample for FIB-SEM imaging as follows: deposition of a protective platinum layer with a 30 kV/3 nA gallium ion beam, followed by coarse trench milling with a 30 kV/30 nA beam, block face polishing with a 30 kV/7 nA beam, and an alternating sequence of FIB milling at 30 kV/3 nA and SEM imaging at 1.5 kV/400 pA electron beam, advancing in 5 nm increments to create isotropic voxels with an X/Y pixel size of 5 nm.

We typically etched fiducial marks into the platinum layer using a 30 Kv/50pa gallium ion beam to form a chevron pattern, which we then infilled with platinum using the SEM at 1.5 kv/5 nA, followed by an additional layer deposited with a 30 kV/1.5 nA gallium ion beam. We collected FIB-SEM images using Inlens and backscatter electron (ESB) detectors, with a pixel dwell time of 10 μs and averaged both images prior to image registration. We then aligned the FIB-SEM images using the Fiji plugin Register Virtual Stack Slices (https://imagej.net/plugins/registervirtual-stack-slices), applying the translation feature extraction and registration model with shrinkage constraint option (Schroeder et al., 2021).

Preparation by conventional chemical fixation of mouse brain samples containing hippocampal CA1 pyramidal neurons, their imaging at 8 x 8 x 20 nm resolution using our Zeiss Crossbeam 540 and availability of data sets are described in the previous study by (Sheu et al., 2022). We also analyzed similar images collected with 5.5 x 5.5 x 15 nm resolution at Janelia Research Campus (Sheu et al., 2022). Preparation by conventional chemical fixation of the mouse liver and skin P7 Skin samples, and their imaging collected with 8 x 8 x 8 nm resolution at Janelia Research Campus are described in (Parlakgül et al., 2022) and OpenOrganelle, HHMI, respectively.

Visual inspection of endosomes and lysosomes was carried out using Neuroglancer (Neuroglancer), the WebGL-based viewer for volumetric data. Cropped regions of interest containing selected endosomes and lysosomes were averaged by 3x in all three dimensions before using them to segment the limiting membranes with the FIJI plugin Labkit (Arzt et al., 2022). The resulting masks were exported into Imaris (Bitplane; Zurich, Switzerland) using the “surfaces” feature of Imaris; movies were created using the ‘animation’ feature of Imaris.

### Image analysis

Data obtained to determine extent of colocalization and ratios between appropriate combinations of fluorescence signals of α-syn-YFP aggregates, mCherry-galectin-3, mCherry-galectin-8, pH sensitive Dextran pHrodo Green 488, pH-insensitive Dextran Alexa Fluor 560, and antibodies specific for EEA1 or LAMP1 were obtained using 3D spinning disc confocal microscopy. Similarly, data obtained to determine the extent of uptake of Transferrin AF647, PFF-AF647 and Dextran AF647 was obtained using live 3D spinning disc confocal microscopy.

Automation to facilitate the extraction of data and analysis (colocalization, ratio, and uptake) was done using a combination of FIJI macros, MATLAB, and Python available upon request.

Colocalization of the fluorescent signals for objects in all experiments carried in the absence of Apilimod treatment was done as follows: (a) identify pixels above background and generate a 3D binary mask (0-1 values) using 3D CME Analysis software (Aguet et al., 2016); z-axis transformed into time. (b) transform time into z-axis and re-scale the binary image to 0-255 values using the segmentation function in the 3D suite of Fiji (Ollion et al., 2013). (c) create 3D binary objects by linking consecutive binary masks along the z-axis using *labelling* within the segmentation function. (d) calculate the logical intersect between each 3D binary mask in one channel with the adjacent 3D binary mask in the second channel using multiplication in Fiji. (e) for each object and for each channel, calculate the extent of colocalization by determining the ratio between the integrated binary values within the logical intersect and the 3D binary object. Colocalized objects had equal or more than 50% volumetric overlap.

The ratio of fluorescence signals between the pH sensitive Dextran pHrodo Green 488 and pH-insensitive Dextran Alexa Fluor 560 was obtained in the absence or presence of Apilimod, as follows. In the absence of Apilimod, calculate the 3D binary mask for the Dextran Alexa Fluor 560 objects following the step (a) just described. (f) In the presence of Apilimod, calculate the 3D binary mask for the Dextran Alexa Fluor 560 objects using LabKit (Arzt et al., 2022); this step was required because the expanded endosomes and lysosomes generated by Apilimod were not properly recognized by 3D-CME. (g) Create the 3D binary mask as described in step (b,c). (h) calculate the integrated fluorescence intensity for each object defined in the 3D binary masks, by using *quantify 3D* within the 3D suite of Fiji after correcting the raw images by the global background. (i) Calculate the ratio of integrated fluorescence intensity between the pH sensitive and pH insensitive dextran. (j) Convert ratio to pH using the pH calibration curve; data presented as a frequency distribution of pH per object, with a binning of 0.5 pH units.

The endocytic uptake of the molecules of interest was calculated from the integrated fluorescence intensity per object determined following (h) and based on the 3D binary mask obtained according to (a-c) without Apilimod or (f,b,c) with Apilimod. (k) The total uptake at a given time point was estimated by adding all the integrated fluorescence intensity values within a given 3D z-stack.

We quantified cell viability by establishing the number of live and dead iNs per imaging field identified by their fluorescence labelling with calcein AM and BOBO-3 imaged using spinning disc confocal microscopy. Data was process using the cell counter plugin in ImageJ/FIJI and statistical analysis was done using the 2-way Anova with Dunnet’s post hoc test in GraphPad Prism 10.0. Data is presented as mean ± standard deviation of the total number and percentage of live and dead cells, with each point corresponding to results from a single imaging field, from a total of 3 biologically independent sets of differentiated iNs, and a total of 40-50 iNs per condition and per experiment counted. The symbol * corresponds to a statistically significant difference of p<0.0001.

**Video 1.** Perforated endolysosome from an iN. The video starts with a series of consecutive single plane images traversing the endolysosome depicted in Fig. 4A, visualized using focused ion beam scanning electron microscopy (FIB-SEM), concluding with a three-dimensional surface depiction of the limiting membrane. This rendering accentuates a single perforation corresponding to a nanopore situated within the limiting membrane.

**Video 2.** Perforated endolysosome from an iN. The video starts with a series of consecutive single plane images traversing the endolysosome depicted in Fig. 4B, visualized using focused ion beam scanning electron microscopy (FIB-SEM), concluding with a three-dimensional surface depiction of the limiting membrane. This rendering accentuates a single perforation corresponding to a nanopore situated within the limiting membrane.

**Video 3.** Intact endolysosome from an iPSC. The video starts with a series of consecutive single plane images traversing the endolysosome depicted in Fig. 4C, visualized using focused ion beam scanning electron microscopy (FIB-SEM), concluding with a three-dimensional surface depiction of the limiting membrane. This rendering accentuates the absence of perforations in the limiting membrane.

## ACKNOWLEDGMENTS

We thank S.C. Harrison for extensive editorial help; and members of the Kirchhausen laboratory for help and encouragement. The research was supported by a National Institute of General Medical Sciences Maximizing Investigators’ Research Award GM130386 and a generous grant from IONIS to T. Kirchhausen. Athul Nair was supported in part by discretionary funds available to T. Kirchhausen. Acquisition of the FIB-SEM microscope was supported by a generous grant from Biogen to T. Kirchhausen, and the high-pressure freeze substitution device was made available by S.C. Harrison.

## The authors declare no competing financial interests

## Author contributions

A. Sanyal and T. Kirchhausen conceptualized and designed the experiments; A. Sanyal carried the cell biological experiments; in vivo pH experiments were jointly done by A. Sanyal and A. Saminathan; A. Sanyal and E. Somerville prepared the samples for FIB-SEM; E. Somerville maintained the FIB-SEM, and also collected and processed the FIB-SEM volumetric data; A. Nair and E. Somerville analyzed the volumetric FIB-SEM data; G. Scanavachi carried out the optical image analysis; A. Oikonomou with supervision from N. S. Hatzakis developed the python based automation tools used for analysis of the optical images; T. Kirchhausen drafted the manuscript and contributed to its final version in close consultation with the co-authors.

